# Antisense transcription and PRC2 repression function in parallel during vernalization

**DOI:** 10.1101/2023.07.07.547987

**Authors:** Mathias Nielsen, Govind Menon, Yusheng Zhao, Eduardo Mateo-Bonmati, Philip Wolff, Shaoli Zhou, Martin Howard, Caroline Dean

## Abstract

Non-coding transcription induces chromatin changes that can mediate environmental responsiveness, but the causes and consequences of these mechanisms are still unclear. Here, we investigate how antisense transcription interfaces with Polycomb Repressive Complex 2 silencing during winter-induced epigenetic regulation of Arabidopsis *FLOWERING LOCUS C* (*FLC*). Through genetic, chromatin, and computational analyses, we show that *FLC* is silenced through pathways that function with different dynamics: an antisense transcription-mediated pathway capable of fast response; and in parallel a slow Polycomb Repressive Complex 2 (PRC2) switching mechanism that maintains each allele in an epigenetically silenced state. Components of both the antisense and PRC2 pathways are regulated by a common transcriptional regulator (NTL8), which accumulates slowly due to reduced growth at low temperatures. The parallel activities of the regulatory steps, which we encapsulate in a mathematical model, creates a flexible system for registering widely fluctuating natural temperature conditions that change year on year, and yet ensure robust epigenetic silencing of *FLC*.

**Significance:** The role of non-coding transcription in establishing and maintaining chromatin states is controversial, mainly because of extensive feedbacks complicating analysis of the relationship between co-transcriptional processing, chromatin state and transcription. This controversy has extended to the role of antisense transcription in the Polycomb-mediated epigenetic silencing of Arabidopsis *FLC*, a key step in the process of vernalization. Here, we show that antisense transcription and PRC2 silence *FLC* in parallel pathways that are affected by growth dynamics and temperature fluctuations. These features explain the varied importance of antisense transcription in cold-induced *FLC* epigenetic silencing seen in various studies using different environmental and growth conditions. The parallel repressive inputs and extensive feedbacks make the mechanism counter-intuitive but provide great flexibility to the plant.

## Introduction

Non-coding transcription has emerged as an important mechanism in environmentally responsive gene regulation. In some cases, non-coding transcription induces chromatin changes that are lost if the environmental signal is removed (1, 2). In other cases, chromatin changes, particularly those involving the Polycomb mark H3K27me3, are epigenetically maintained, thus providing a memory of the inductive signal. One well-characterised example of the latter is the winter-induced epigenetic silencing of the Arabidopsis floral repressor gene, *FLC* (3, 4). This underpins the vernalization process, the acceleration of flowering by winter exposure. The process includes early induction of a series of antisense transcripts, called *COOLAIR* (5); a slow epigenetic switch from an active chromatin environment (marked by H3K36me3) to a silenced chromatin state (marked by H3K27me3) at an internal three nucleosome region (6); and spreading of the H3K27me3 Polycomb silencing over the whole locus (7, 8). The switching mechanism involves canonical Polycomb Repressive Complex 2 (PRC2) and Arabidopsis PRC2 accessory proteins VIN3 and VRN5. VIN3 is slowly induced by cold exposure (9), interacts with PRC2 at the nucleation region downstream of the *FLC* transcription start site, and has a functionally important head-to-tail polymerization domain (10).

The timing of early antisense transcription and later VIN3 expression led to the view that antisense transcription was a prerequisite for PRC2 silencing. Consistent with this, single-molecule FISH experiments revealed that *COOLAIR* expression was mutually exclusive with *FLC* sense transcription at each allele (11), and reduced transcription was necessary to maintain PRC2 silencing. This sequence of events was initially tested through T-DNA insertions into the *COOLAIR* promoter. These had little effect on long-term vernalization (12). Similarly, *FLC* silencing was unaffected in studies using a CRISPR deletion of the *COOLAIR* promoter or mutation of *CBF* factors, known to facilitate cold-induction of *COOLAIR (13)*. However, replacement of *COOLAIR* 5’ sequences (TEX1 line) attenuated *FLC* transcriptional silencing and disrupted the co-ordinated changes in H3K36me3 and H3K27me3 occurring at the *FLC* nucleation region (14).

*COOLAIR* had much stronger effects in experiments analysing *FLC* silencing in natural field conditions. *COOLAIR* expression was strongly induced on the first freezing night of autumn (15, 16), a result re-created in controlled environment cabinets (15). In these experiments, one freezing night was sufficient to induce *COOLAIR*, but several freezing nights were required to silence *FLC*, with silencing attenuated by disruption of antisense transcription. These data are reminiscent of many *S. cerevisiae* loci, where non-coding transcription plays an important role in environmental responsiveness (1, 17, 18). However, extensive feedback mechanisms between chromatin, transcription and cotranscriptional processes make function of non-coding transcription difficult to elucidate. In particular, buffering between transcription and RNA stability leads to changed transcriptional dynamics with no change in steady state RNA (2).

To clarify the regulatory mechanism at Arabidopsis *FLC*, we have undertaken a series of genetic, molecular and computational analyses to investigate the role of *COOLAIR* in cold-induced *FLC* silencing. Here, we show that *FLC* is silenced through parallel pathways. On a fast timescale *COOLAIR* limits sense transcription, and this is associated with reduction in levels of the active histone mark H3K36me3. This mechanism involves disruption of a 5’ -3’ *FLC* gene loop. On a slower timescale, PRC2 silencing switches each allele from an epigenetically ON to an OFF state. This involves nucleation of H3K27me3 and subsequent spreading over the locus during active cell cycle. The nucleated and spread states differentially influence *FLC* transcription, which is still modulated by antisense transcription. Components of both pathways are also regulated by their common transcriptional regulator (NTL8) (19), which accumulates based on reduced dilution dependent on growth dynamics in the different cold phases. We integrate these parallel regulatory activities into a mathematical model that predicts *FLC* chromatin dynamics in different conditions. We argue that parallel activities converging onto a common target provides great flexibility in gene regulation, providing responsiveness to a wide variety of conditions. The extensive similarities between how antisense transcription modulates *FLC* and how it alters sense transcription dynamics in yeast (2) suggest these findings will be generally important.

## Results

### *COOLAIR* rather than PRC2 nucleation is the major contributor to *FLC* repression in *ntl8-D3*

Two independent genetic screens in different genotypes had identified dominant mutations that revealed NTL8 regulated *VIN3* and *COO-LAIR* (15, 19). Ectopic *COOLAIR* expression led to very low *FLC* levels in warm grown plants (15). We confirmed that *VIN3* and *COOLAIR* are both misregulated in the dominant mutant *ntl8-D3* (Fig. 1A) (15, 19) and then used it to genetically activate both pathways simultaneously, independently of cold. *FLC* transcriptional output, histone modifications and chromatin topology were analysed. Paralleling cold effects on wild-type plants the ec-topic *COOLAIR* expression in *ntl8-D3* resulted in a clear decrease in H3K36me3 as compared to Col*FRI* at the *FLC* transcription start site (TSS) and over the gene body (Fig. 1B) (20, 21). The high *COOLAIR* transcription in *ntl8-D3* led to accumulation of H3K36me3 at the *COOLAIR* promoter (Fig. 1B), matching the cold-induced transient increase of H3K36me3 in Col*FRI* at the same position (20). The decrease in H3K36me3 was not accompanied by an increase in H3K27me3 as observed during vernalization (Fig. 1C). Like-wise, H2Aub, another histone modification that accumulates at *FLC* during early vernalization (22) did not accumulate ectopically in *ntl8-D3* (Fig. 1D). The lack of accumulation of H3K27me3 and H2AUb in *ntl8-D3* compared to Col*FRI* in the absence of cold supports the view that VIN3 expression itself is not sufficient to cause Polycomb mediated silencing of *FLC (23)*. These data indicate that antisense-mediated suppression rather than VIN3-mediated nucleation of H3K27me3 is the major factor causing *FLC* repression in *ntl8-D3*. The same conclusion was made after the repression of sense *FLC* transcription in *ntl8-D3* was found to be suppressed by transgenes interfering with *COOLAIR* transcription (15).

**Figure 1.**
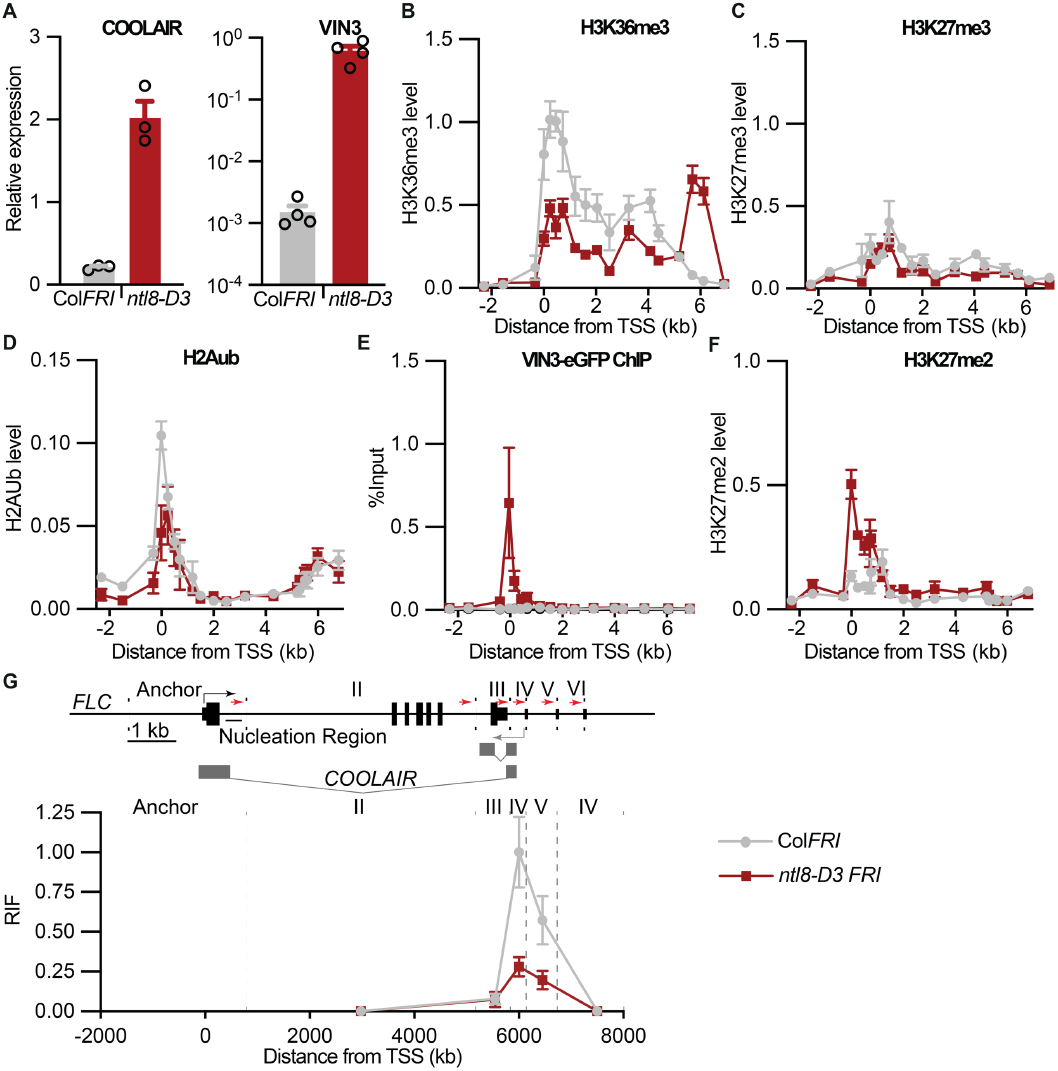
*ntl8-D3* mimics cold exposure, except for the accumulation of H3K27me3. **(A)** Expression of total *COOLAIR* and *VIN3* in *ntl8-D3 FRI* and in Col*FRI* in non-vernalized conditions (NV). Data are presented as the mean ± s.e.m. Each open circle represents a biological replicate. (**B-D**) Enrichment of **(B)** H3K36me3, **(C)** H3K27me3, and **(D)** H2AUb across *FLC* measured by ChIP in wild-type Col*FRI* and *ntl8-D3* at NV conditions. H3K36me3 data are shown relative to H3, and actin. H3K27me3 data are shown relative to H3, and STM. H2Aub data are shown relative to H3. Error bars represent s.e.m (n ≥ 3 biological replicates). **(E)** VIN3-eGFP ChIP-qPCR enrichment at *FLC* at NV. Data are shown as the percentage input. Non-transgenic Col*FRI* plants were used as a negative control sample. Error bars represent s.e.m (n = 3 biological replicates). **(F)** Enrichment of H3K27me2 across *FLC* measured by ChIP in wild-type Col*FRI* and *ntl8-D3* at NV conditions. Data are expressed relative to H3. Error bars represent s.e.m (n = 3 biological replicates). **(G)** Quantitative 3C-qPCR over the *FLC* locus in 10-day-old Col*FRI* and *ntl8-D3 FRI* seedlings. A schematic representation of the *FLC* locus is shown above. BamHI and BglII restriction sites are indicated with dotted lines, and the respective regions are numbered with Roman numerals. Red arrows indicate the location of the primers used for 3C-qPCR. The region around the *FLC* transcription start site was used as the anchor region in the 3C analysis. The data below shows the relative interaction frequencies (RIF).

### Ectopically expressed VIN3 localises to *FLC* but fails to induce H3K27me3 nucleation

To understand what prevents the accumulation of H3K27me3 in *ntl8-D3* despite ectopic VIN3 expression, we tested if other epigenetic factors are misexpressed in *ntl8-D3*. Only one of the tested genes changed slightly in expression (SI Fig. 1). We then analysed the association of VIN3 at the nucleation region in *ntl8-D3*. Despite the lack of H3K27me3 accumulation in *ntl8-D3*, we found VIN3-eGFP accumulated at the *FLC* nucleation region in warm conditions, mimicking the accumulation during vernalization (Fig. 1E). Thus, VIN3 accumulation at the nucleation region does not result in stable nucleation of H3K27me3. To distinguish VIN3 intrinsic binding to the *FLC* nucleation region, independently of *COOLAIR* transcriptional induction, we expressed VIN3-eGFP under the promoter of VRN5 (SI Fig 2A). This resulted in expression levels in non-vernalized plants that paralleled VIN3 induction after six weeks cold (6WT0) (SI Fig. 2B). In this line VIN3-eGFP was enriched at the *FLC* locus in NV conditions (SI Fig. 2C), showing VIN3 can remain associated with the nucleation region even when *FLC* is strongly expressed. We found that VIN3 association in *ntl8-D3* led to H3K27me2 enrichment despite no accumulation of H3K27me3 (Fig. 1F). Thus, cold-induced features, possibly influencing residence time, are required to enable VIN3 functionality to deliver H3K27me3 to the nucleation region.

**Figure 2.**
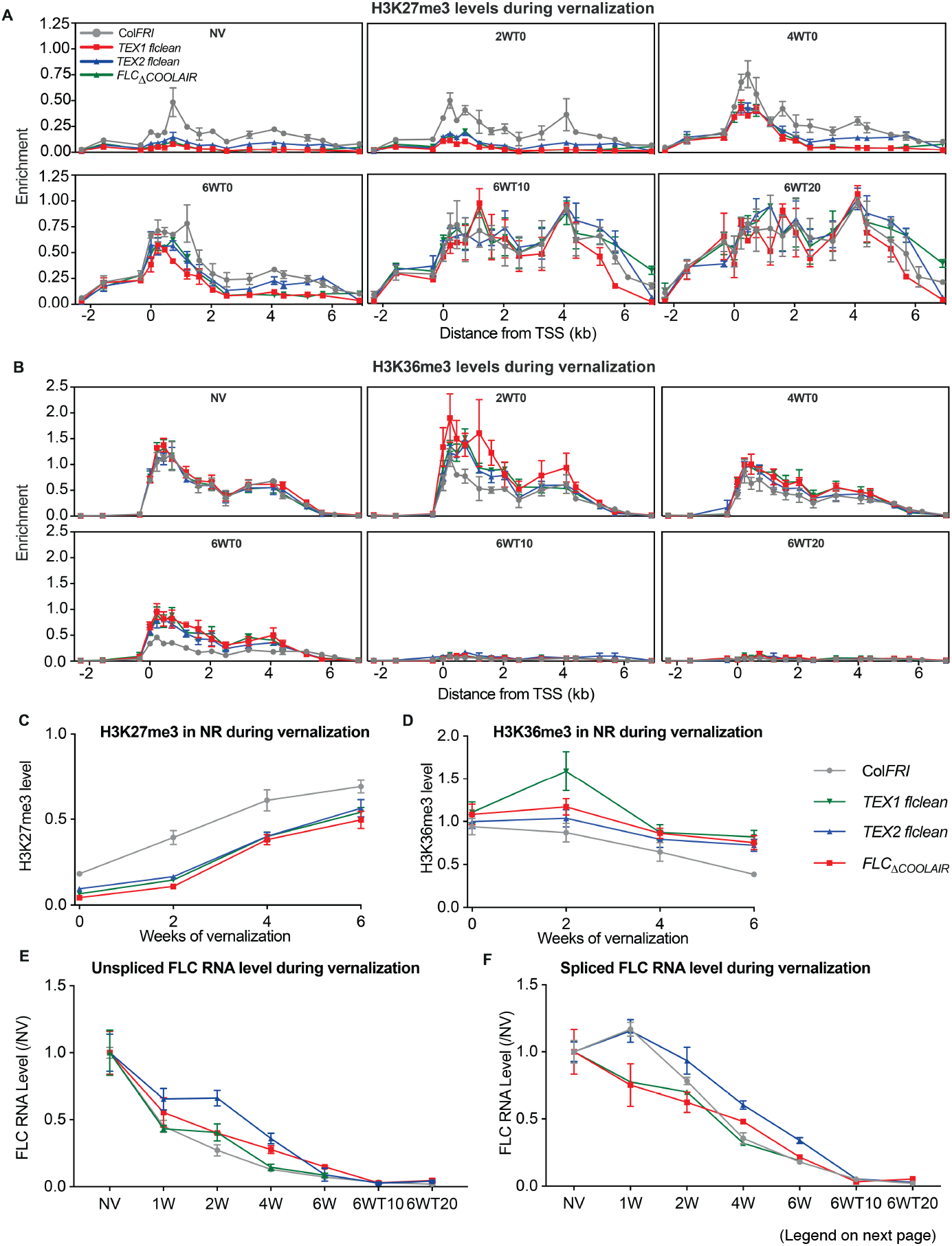
Cold-induced chromatin and RNA dynamics in *COOLAIR* defective lines. **(A-B)** Enrichment of H3K27me3 **(A)** and H3K36me3 **(B)** across *FLC* measured by ChIP in wild-type Col*FRI* and the three defective *COOLAIR* lines, *TEX1, TEX2, and FLC*_*≥COOLAIR*_, before, during, and after vernalization. Data are expressed relative to H3 and *STM*. Error bars represent s.e.m. (n = 3 biological replicates). **(C-D)** Average levels of H3K27me3 **(C)** and H3K36me3 **(D)** in the nucleation region during vernalization. The averages were calculated by averaging the ChIP enrichment over three primers in the *FLC* nucleation region during vernalization in Col*FRI* and each of the defective *COOLAIR* lines. **(E-F)** *FLC* expression during a vernalization time course in Col*FRI* and the three defective *COOLAIR* lines, Unspliced **(E)** and spliced RNA **(F)**, was measured and is shown relative to *UBC* and NV levels. Error bars represent s.e.m. (n = 3 biological replicates).

### Ectopic induction of *COOLAIR* correlates with chromatin topology changes

A cold-induced feature at *FLC* is disruption of a gene loop conformation that links the transcription start site (TSS) and the transcription termination site (24). In *ntl8-D3*, we found that the gene loop was ectopically disrupted, mimicking vernalization (Fig. 1G). This suggests that gene loop disruption is tightly linked with antisense-mediated reduction in *FLC* transcription. We also found that the TEX2.0 transgene, where a *nos* terminator promotes early *COOLAIR* termination, reduces gene loop formation (SI Fig. 3), consistent with earlier reports using a similar, but not identical transgene (25). This result suggests a role for the activity of the antisense promoter/TSS, rather than antisense transcription per se, as being important for gene loop disruption.

### Disrupting *COOLAIR* transcription perturbs H3K27me3 dynamics before and during cold, but not post-cold H3K27me3 levels

We further investigated the fact that ectopic expression of antisense transcription is enough to cause lower H3K36me3 around the *FLC* sense TSS and in the gene body, even in the absence of cold. Antisense transcription could lower H3K36me3 levels, either through direct removal mediated by antisense transcription or indirectly by limiting sense transcription, thus preventing the cotranscriptional addition of H3K36 methylation. To dissect the interplay of H3K36me3 and H3K27me3, we studied the dynamic changes in these modifications using a vernalization time course.

Our previous analyses of TEX transgenes was in an *flc-2* background, where part of the endogenous *FLC* genomic sequence remains (4). This limited the regions where the chromatin modifications on the transgene could be studied (14). To overcome this limitation, we generated a FRI +, *FLC* null (*flclean*) where the entire *FLC* genomic sequence had been deleted using CRISPR (SI Fig. 4) and introduced the previously described TEX1.0 (replacement of the *COOLAIR* promoter) and TEX2.0 (insertion of a *nos* terminator to truncate *COOLAIR* transcription) transgenes. We also included a *FRI FLC*_*ΔCOOLAIR*_ CRISPR line, which deletes the *COOLAIR* promoter at the endogenous locus (26). Using these multiple defective *COOLAIR* lines and respective controls, we undertook a detailed time course of histone modifications during vernalization, including multiple time points post-cold (Fig. 2A, B).

The rate of accumulation of H3K27me3 during cold exposure was not reduced in the *COOLAIR* defective genotypes compared to the wild-type, and at some timepoints was even accelerated (Fig. 2C and SI. Fig 5A,B), consistent with our previous data (14). By 6WT0, wild-type and *COOLAIR* defective genotypes show similar H3K27me3 levels in the nucleation region (Fig. 2A and SI Fig 5C). However, there were clear differences in the starting levels of H3K27me3 – being significantly lower in the nucleation region in all defective *COOLAIR* genotypes (Fig. 2A and SI Fig 5D). The relative levels of H3K27me3 matched *FLC* RNA in the different *COOLAIR* genotypes before cold exposure (SI Fig. 5E), supporting a role for antisense transcription in establishment of the initial *FLC* chromatin state (27). The similar trend in TEX1, TEX2 and *FLC*_ΔCOOLAIR_ argues against this being a transgene variability effect. We interpret the H3K27me3 level in Col*FRI* before cold as representing a fraction of *FLC* alleles that have switched to a stable Polycomb silenced state. Thus, higher H3K27me3 levels in Col*FRI* compared to the *COOLAIR* defective genotypes may reflect the antisense role in developmentally regulated PRC2 silencing of *FLC (28)*. After cold there was no significant difference in H3K27me3 levels in the nucleation region between Col*FRI* and any of the *COOLAIR* defective genotypes (Fig. 2A and SI Fig 5C). Spreading of H3K27me3 was also unaffected in the *COOLAIR* defective genotypes, as seen from the similar levels in the gene body at 6WT10 and 6WT20 (Fig. 2A). Overall, we find that H3K27me3 dynamics before and during cold are perturbed by *COOLAIR*, but that post-cold H3K27me3 levels are not.

### Disrupting *COOLAIR* transcription attenuates H3K36me3 removal during vernalization

H3K36me3 levels were similar in all genotypes before vernalization (Fig. 2B and SI Fig 5F) but decreased at different rates during cold exposure (Fig. 2D). This is in contrast to the clear NV differences in H3K27me3 levels. However, this is consistent with the NV H3K27me3 levels coming from a small fraction of silenced alleles, while the majority of alleles are transcriptionally active and contribute to the observed H3K36me3 levels, a scenario that generates bigger fold changes in H3K27me3 than in H3K36me3, as we observe (SI Fig 5G). H3K36me3 levels reduced more slowly in all defective *COOLAIR* genotypes at 6WT0 (Fig. 2B and SI Fig 5G), but after 2 weeks cold H3K36me3 levels increased in the gene body compared to NV (Fig. 2B). There were no differences in H3K36me3 levels between *COOLAIR* defective genotypes and wild-type Col-*FRI* after transfer back to warm (Fig. 2B). In *ntl8-D3*, where *VIN3* and *COOLAIR* are both overexpressed, faster reduction of H3K36me3 in the cold was observed, while H3K27me3 was barely affected (Fig. S6 A-D, Fig. S9B,C). Together our results demonstrate that the Polycomb pathway is effective enough to completely silence the *FLC* locus, despite either an ineffective or hyperactive antisense pathway. The antisense-mediated pathway mediates not only the removal of H3K36me3 but also H3K4me1 through the activity of the demethylase complex FLD-LD-SDG26 (27). H3K4me1, like H3K36me3, has been shown to be added co-transcriptionally in plants (29). Consistently, we found that in the *COOLAIR* defective lines, H3K4me1 reduction during vernalization was attenuated (SI Fig S7) showing the same trend as H3K36me3, including the increase at 2WT0. Overall, we find that *COOLAIR* defective genotypes have reduced rates of H3K36me3 removal, but after cold, any differences in H3K36me3 levels disappear. The relative changes in unspliced *FLC* RNA levels (often taken to reflect nascent transcripts) did not match the corresponding H3K36me3 levels in the *COOLAIR* defective genotypes (Fig. 2E). The relative effects of *COOLAIR* modifications on spliced *FLC* levels were also different to those of unspliced *FLC* RNA (Fig. 2F). This suggests a similar interconnected mechanism linking chromatin modification to transcript stability as found in yeast, with unspliced and spliced transcripts affected in different ways (2).

### H3K27me3 accumulation is not necessary for *COOLAIR-* mediated transcriptional downregulation

A mutation in the core PRC2 component Su(z)12 (VRN2) only partially disrupted *FLC* repression (SI Fig. S8A) (6) while H3K36me3 fold reduction at *FLC* during the cold was hardly changed (SI Fig. S8B), despite accumulation of H3K27me3 being abolished (6). Thus, H3K36me3 reduction and *FLC* RNA downregulation do not appear to rely on H3K27me3 nucleation. Analysis of a *vrn5-TEX1*.*0* combination, defective in H3K27me3 accumulation and *COOLAIR*, had shown that the H3K36me3 reduction seen in an H3K27me3 nucleation mutant is mediated by *COOLAIR (14)*. To examine this aspect further, we analysed changes in the two modifications – H3K36me3 and H3K27me3 – in fluctuating cold conditions, where we see the clearest indication of *COOLAIR* transcription regulating *FLC* expression (15). Under these conditions, *COOLAIR* was highly upregulated, causing significant downregulation of *FLC* sense transcript (15). While we have previously shown that full-length *COOLAIR* transcription is essential for the *FLC* downregulation in these conditions (15), a role for H3K27me3 nucleation had not been investigated. Here we analysed H3K36me3 and H3K27me3 levels in Col*FRI* at 2WT0, under three different cold conditions (as in (15)), constant 5°C (CC, Constant Cold), mild 3°C -9°C (FM, Fluctuating Mild), and strong fluctuating conditions -1°C - 12°C (FS, Fluctuating Strong). Zhao et al., 2021 showed that *COOLAIR* upregulation and *FLC* downregulation were greatest in the FS condition. Therefore, we would expect H3K36me3 to show the largest changes in this condition, and indeed this is seen in our data (Fig. 3A, SI Fig. S9E). Both mild fluctuating and constant cold result in smaller changes (Fig. 3A, SI Fig. S9E). In contrast, H3K27me3 accumulation showed little difference between the different conditions (Fig. 3B and SI Fig. S9D), indicating that H3K27me3 is not the major contributor to the enhanced downregulation under strongly fluctuating conditions. The lack of difference in H3K27me3 accumulation, despite the relatively large change in antisense expression, further highlights the parallel and almost independent nature of these *FLC* repression pathways. To further confirm that *COOLAIR* transcription is necessary for the changes in H3K36me3 under fluctuating conditions, we subjected the TEX1.0 defective *COOLAIR* line to these conditions. As expected, the reduction in H3K36me3 was also attenuated relative to Col*FRI* (Fig. 3C, D) (30). Interestingly, the slight increase in H3K36me3 at 2WT0 observed in constant cold was also recapitulated in the FM conditions. The reduction under FS conditions was significantly attenuated in TEX1.0 (SI Fig. S9F). Overall, we find that *COOLAIR*-mediated transcriptional repression does not strongly depend on H3K27me3 nucleation, supporting our earlier results in *ntl8-D3*. The contrasting dynamics of H3K36me3 and H3K27me3 under 2 weeks of fluctuating conditions further highlight the fast-response capability of the antisense-mediated repression. In Col*FRI*, the striking changes in nucleation region H3K36me3 after only 2 weeks of FS conditions (Fig. 3A,C) are comparable to H3K36me3 changes after 6 weeks in constant cold (Fig. 2B, SI Fig. 5G).

**Figure 3.**
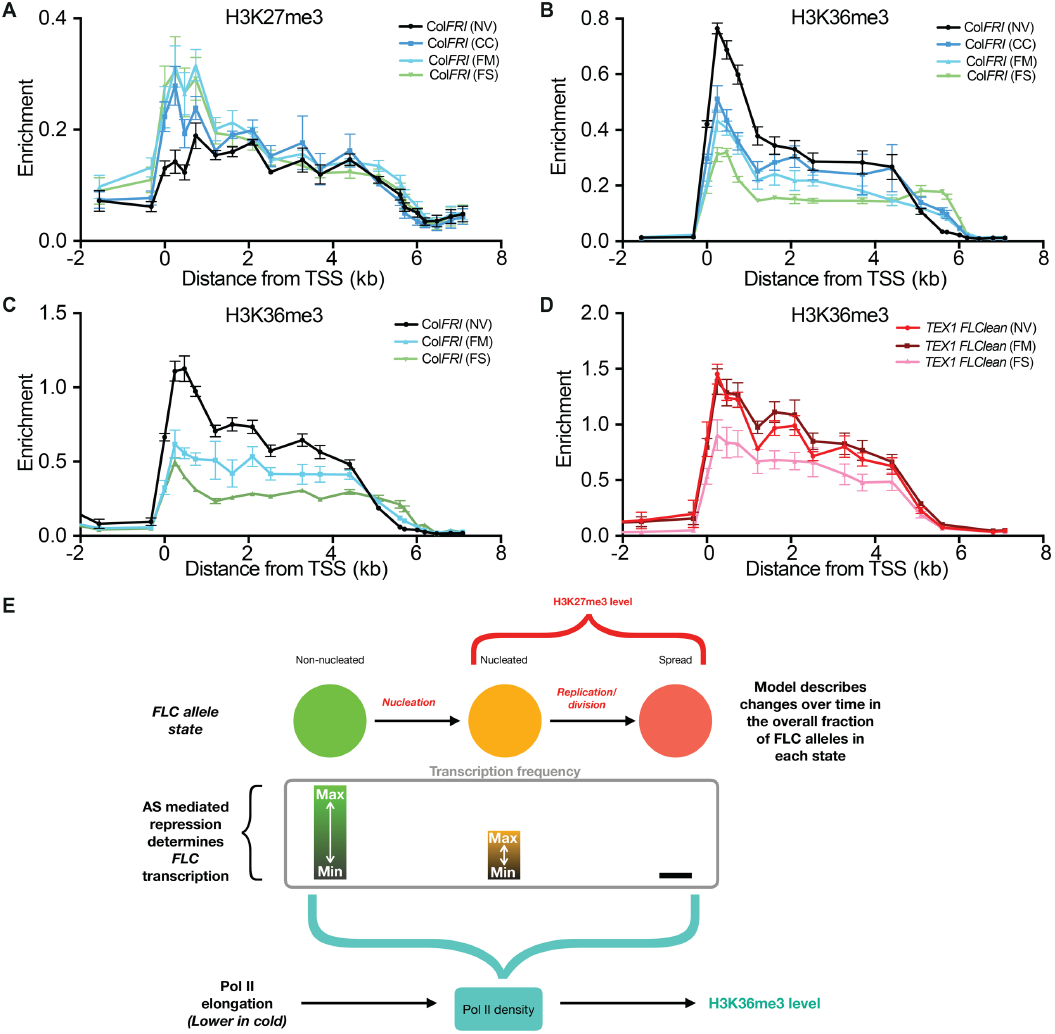
Fluctuating cold and mathematical modelling of antisense role in histone modification dynamics. **(A-B)** Changes in H3K36me3 **(A)** and H3K27me3 **(B)** at *FLC* after two weeks of Constant Cold (CC), Fluctuating Mild (FM), or Fluctuating Strong (FS) conditions, measured by ChIP-qPCR. Data are expressed relative to H3. Error bars represent s.e.m. (n = 3 biological replicates). **(C-D)** Comparing changes in H3K36me3 at *FLC* between Col*FR*I **(C)** and *TEX1 flclean* **(D)** after two weeks of FM or FS conditions, measured by ChIP-qPCR. Data are expressed relative to H3 and *STM*. Error bars represent s.e.m. (n = 3 biological replicates). **(E)** Schematic of the mathematical model showing core components: PRC2 mediated silenced states (non-nucleated, nucleated and spread) at individual *FLC* alleles, antisense transcription mediated repression of *FLC* transcription, and the contribution of these components to the average population level H3K36me3 coverage at the *FLC* locus.

### Mathematical modelling of *FLC* regulation reconciles the different effects of antisense transcription on chromatin state

The repressive dynamics delivered by both pathways are difficult to dissect quantitatively purely through molecular experiments. We, therefore, turned to mathematical modelling to see how the observed behaviour in *COOLAIR* defective mutants could be reconciled with our existing understanding of *FLC* repression in the cold. We have previously developed and experimentally validated a mathematical modelling framework describing dynamically changing fractions of active/silenced *FLC* alleles and their associated histone modifications (31, 32). Here we built on this framework to develop a new model, incorporating an antisense-mediated silencing pathway. A schematic of the model developed here is shown in Fig. 3E (details in Supplementary Information). The model was built on the basis of our main conclusion from the above data: namely that two pathways work in parallel to silence *FLC*, antisense transcription and PRC2 nucleation. We then interrogated the model to see whether it was capable of quantitatively reproducing histone modification dynamics in Col*FRI* and the various mutants.

The effect of the antisense-mediated pathway on sense transcription was modelled implicitly as a cold-dependent graded modulator of sense initiation/transition to productive elongation. This is consistent with high antisense transcription in *ntl8-d* mutants causing low levels of *FLC* transcription, independently of H3K27me3 nucleation. This is also consistent with previous data showing that sense and antisense transcription at *FLC* are anti-correlated in Col*FRI*, both in warm (33) and in cold conditions (11). This may be through a mutual exclusivity model for the *FLC* locus, similar to that reported for the *CBF1-SVALKA* locus, where full-length sense transcription is inhibited by antisense transcription (34). Another key aspect of the model is the cotranscriptional delivery of the H3K36me3 modifications. Changes in Pol II elongation behaviour can affect the H3K36me3 profile across mammalian genes (35), with slower Pol II elongation allowing a larger window of opportunity for adding H3K36me3 at any given location. Any changes in transcription at *FLC* may be expected to produce corresponding changes in H3K36me3. However, to explain the *increase* in gene body H3K36me3 observed in the defective *COOLAIR* lines, specifically at 2WT0 compared to NV, despite the lack of any increase in transcriptional output over that time period (Fig. 2E), the model includes a cold-induced reduction in Pol II elongation in this region, resulting in a longer dwell time. The SDG8 H3K36 methyltransferase, which we have shown co-transcriptionally associates with RNA PolII (20), is likely mediating these H3K36me3 changes.

We also incorporate the PRC2 pathway and how it silences *FLC* through H3K27me3 accumulation during vernalization. In cells which can have active or non-active cell cycles, we consider that *FLC* alleles can be in three different states – non-nucleated (without H3K27me3 nucleation), nucleated, and spread states, with the latter only attained in active cycling cells, as previously introduced (30). To generate reasonable fits to our data, particularly the higher levels of H3K36me3 observed during cold in *COOLAIR* defective mutants, we found that an extension to our previous models was needed, where we allow for different levels of *FLC* transcription in the three states: highest in non-nucleated, much lower in nucleated and even lower in spread. Satisfactory fits also necessitated that the non-nucleated and nucleated states be capable of further downregulation by antisense transcription. Consistent with the possibility of some limited transcriptional activity in the nucleated state, we, therefore, allowed for potential co-existence of H3K27me3 and H3K36me3 on the same nucleosome. We then fitted the model to capture the qualitative changes in H3K36me3, H3K27me3, and transcription (sense and antisense) observed in the cold for Col*FRI* and the defective *COOLAIR* lines. We found that this model could capture all the qualitative features of the data observed in Col*FRI* and the defective *COOLAIR* lines (Fig. 4A-D), including both the increase of H3K36me3 at 2WT0 and the subsequent significantly slower reduction in H3K36me3 in the latter (Fig. 4A,B), as well as the reduction of H3K36me3 in the post-cold seen in all the lines (Fig.4A,B).

**Figure 4.**
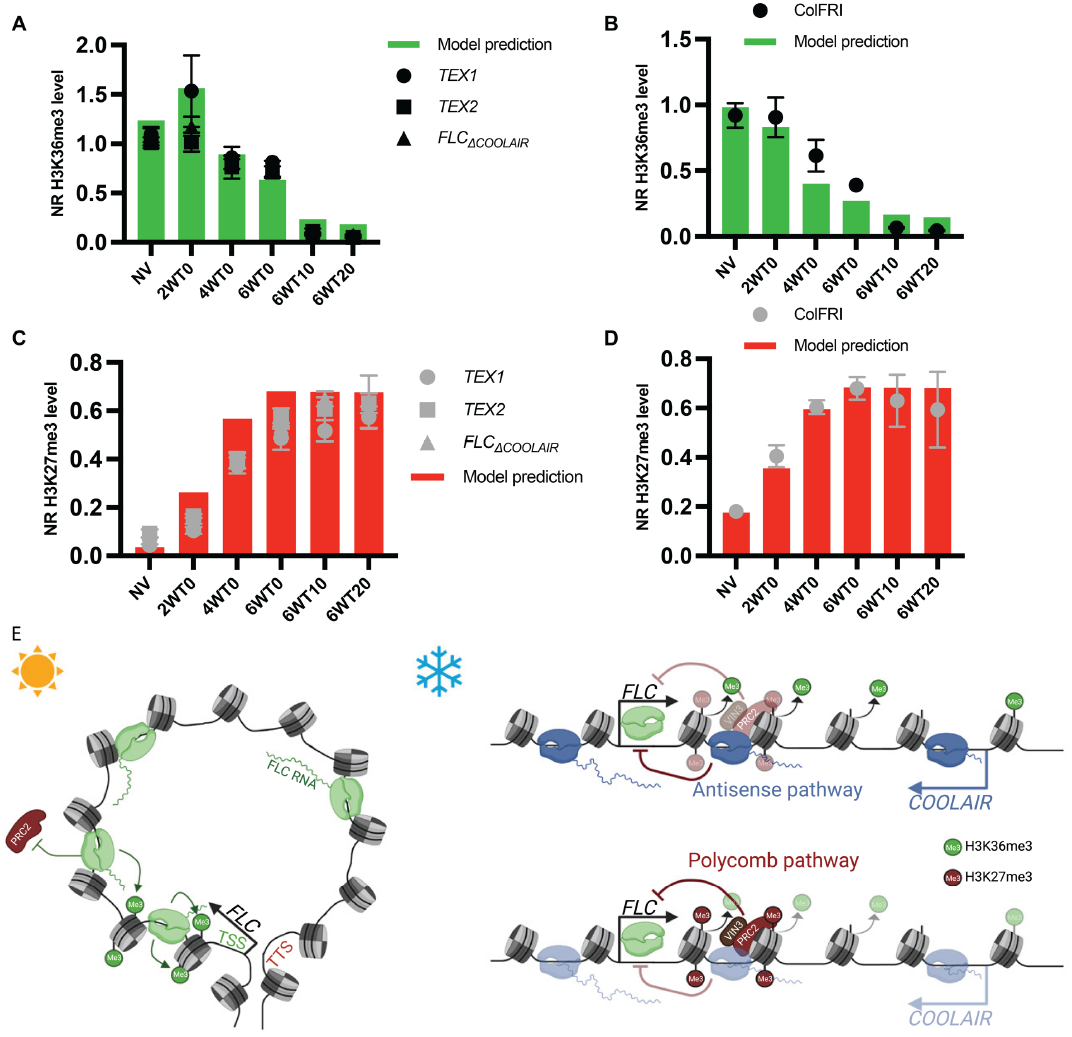
Model predictions of the impact of vernalization mutants on histone dynamics and schematic representation of parallel pathways that repress *FLC* expression. **(A-D)** Mathematical model predicted levels of H3K36me3 **(A**,**B)** and H3K27me3 **(C**,**D)** over a constant cold time course in a defective *COOLAIR* mutant **(A**,**C)** and the wild type, Col*FRI* **(B**,**D)**. The predictions are compared to the ChIP-qPCR time course data for the different genotypes presented in Fig. 2(A,B). **(E)** Model for the parallel pathways that repress *FLC*. In the warm *FLC* forms a gene-loop conformation, which mediates a high expression state of *FLC*. The high expression state is marked by high levels of H3K36me3 around the *FLC* TSS. After cold exposure, the repressive pathways are activated: (1) The antisense-mediated pathway leading to disruption of the gene-loop and removal of H3K36me3 from the TSS of *FLC*. (2) The Polycomb pathway leading to deposition of H3K27me3 and repression of *FLC* transcription. The two pathways work in parallel rather than through a linear sequence of causation to give the final *FLC* expression outcome during vernalization.

We then tested whether the model could capture our previous datasets by simulating other mutants that affect *FLC* silencing in the cold (see previously published data (6)), including an H3K27me3 nucleation mutant (e.g., *vrn2, vin3*), a spreading mutant (e.g., *lhp1,clf)*. In all cases, the simulation outputs from the model are qualitatively consistent with data, including the post-cold behaviour of the two histone modifications (SI Fig. S10A-D). Interestingly, in addition to recapitulating the behaviour captured by our previous models, the new model is able to capture the reduction of H3K36me3 in the post-cold seen in Col*FRI* - a trend that could not be previously captured (Fig. 4B). This is because the new model allows for higher levels of transcription (and consequently higher H3K36me3) in a nucleated state relative to a spread state.

The model predicts that antisense transcription limits H3K36me3 through a graded, analog reduction in *FLC* transcription rather than by directly mediating H3K36me3 removal. The increased H3K36me3 at 2WT0 in *COOLAIR* defective lines, is predicted to arise from a combination of higher *FLC* sense transcription (since antisense mediated repression is disrupted) and cold-induced slowing of Pol II elongation, resulting in a longer dwell time. In a second slower response chromatin pathway involving Polycomb Repressive Complex 2 (PRC2), each allele progressively switches from a non-nucleated to H3K27me3 nucleated state during the cold and then to a spread H3K27me3 state during post-cold growth. The model indicates that intermediate levels of *FLC* transcription in the nucleated state, which can be further downregulated by antisense transcription, can explain how clear differences in H3K36me3 between defective *COOLAIR* lines and the wild-type can emerge in the cold yet subsequently disappear during growth after transfer to warm conditions. In these conditions, all the nucleated *FLC* alleles convert to the H3K27me3 spread state due to an active cell cycle (6), regardless of H3K36me3 levels and any residual expression. Hence, in the context of vernalization, the *COOLAIR* repressive pathway is most important during rather than after cold.

## Discussion

Focused dissection of the mechanism underlying winter-induced *FLC* silencing has established a role for antisense transcription and PRC2 activity in registration of long-term exposure to noisy environmental signals (5, 15, 36, 37). However, the complexity of the mechanism, and its sensitivity to variable temperature and growth parameters, has led to different studies questioning the importance of the antisense transcription in cold-induced *FLC* silencing. Here, using a combination of experiments and mathematical modelling, we have elucidated the role of antisense transcription and PRC2 activity as parallel pathways, both leading to *FLC* silencing (Fig. 4B). The antisense-mediated pathway involves the *FLC* gene loop and represses *FLC* transcription. Its multiple effects on H3K36me3 in the 5’ region of *FLC* required modelling to deconvolve fully. In wild-type, lower transcription leads to H3K36me3 reduction, but this is partly hidden in defective *COOLAIR* lines in cold through predicted increase in H3K36me3 from lower RNA PolII elongation over the 5’ end of *FLC*. The slow PRC2 switch at each *FLC* allele from a non-nucleated to a H3K27me3 nucleated and then spread state is associated with decreasing frequencies of *FLC* transcription, consistent with previous findings of the relationship of H3K27me3 and *FLC* transcription (38). Both antisense-mediated and PRC2 pathways are affected by the common transcriptional regulator NTL8, which accumulates slowly and variably, dependent on reduced dilution by slower growth at low temperatures (19). Our observation that in *ntl8-D3* both *COOLAIR* and *VIN3* are ectopically expressed, yet the H3K27me2 modification but not H3K27me3 accumulates, implies a requirement for other cold-induced factors for vernalization (39, 40).

The antisense-induced chromatin changes are not directly reflected in steady-state unspliced and spliced *FLC* levels, similar to the situation in yeast (2). Which RNA stability mechanisms are involved remain to be determined, but m6A methylation has been shown to regulate *FLC* regulation (41– 43) and is enriched in the *FLC* 3’ UTR, replaced in the TEX1.0 transgene. This disconnect between chromatin dynamics and steady-state RNA levels is likely to go a long way to explaining the controversy as to the role of antisense transcription in the Polycomb-mediated epigenetic silencing of Arabidopsis *FLC* and, potentially, more generally, to the role of non-coding transcription. In addition, the effective combination of parallel pathways hides effects of mutations after saturating induction eg. mutations in CBF-binding factors (13). Another debate has been over the use of transgenes to modulate *COOLAIR* expression (13), but the use here of *FLC*_*≥COOLAIR*_ and *ntl8-D3* for under/overexpression of *COOLAIR* rules this out. However, future studies will need to attempt to generate a fully antisense null genotype, since all defective *COOLAIR* genotypes so far produced still contain cryptic antisense promoters, which become more active when the endogenous *COOLAIR* promoter is mutated/deleted (15). The difficulty of completely removing antisense transcription is also seen in other systems (44) and suggests transcription initiation from open chromatin regions rather than specific promoter elements. Such a line would not only help elucidate the role of *COOLAIR* in the cold-induced silencing of *FLC*, but also in the starting *FLC* expression upon germination, a key determinant of natural variation underpinning adaptation (16).

This study points to similarities between *FLC* silencing and antisense-mediated chromatin modulation of sense transcription dynamics in yeast (2), where antisense expression has been associated with genes that switch off (1). Future work will address the evolutionary parallels and conservation of a mechanism enabling rapid transcriptional changes and switches to epigenetic silencing in response to noisy environmental cues.

## Materials and Methods

Provided as Supplementary Information

## Acknowledgements

The authors would like to thank members of the Dean lab for thoughtful inputs and Shuqin Chen for excellent technical assistance. This work was funded by European Research Council Advanced Grant (EPISWITCH, 833254), Wellcome Trust (210654/Z/18/Z), and a Royal Society Professorship (RP\R1\180002) to CD, and BBSRC Institute Strategic Programmes (BB/J004588/1 and BB/P013511/1) and EPSRC/BBSRC Physics of Life grant (EP/T00214X/1) to MH and CD.

## Notes

### Competing Interest Statement

The authors have declared no competing interest.

