## Supporting information for "Antisense transcription and PRC2 repression function in parallel during vernalization"

bioRxiv

### Supporting Information for
#### Antisense transcription and PRC2 repression function in parallel during vernalization

Mathias Nielsen†, Govind Menon†, Yusheng Zhao, Eduardo Mateo-Bonmati, Philip Wolff, Shaoli Zhou, Martin Howard* & Caroline Dean*

*Corresponding authors Caroline Dean and Martin Howard

**This PDF file includes:**

Supporting text
Figures S1 to S10
Table S1
Legend for Dataset S1
SI References

### SI Materials and Methods

#### Plant materials

All mutant and transgenic lines were in the $FRI^{sf2}$ background. Mutant alleles were described previously: *ntl8-D3 FRI* (1), and $FLC_{\Delta COOLAIR}$ (2). For this study, $FLC_{\Delta COOLAIR}$ was introduced into Columbia with an active *FRIGIDA* allele (3). The TEX1.0 and TEX2.0 constructs were described previously (1, 4), and for this study, the constructs were introduced into *FLClean* by floral dipping. *FLClean* was created by CRISPR-CAS9. The CRISPR guide-RNA sequences were: sgRNA_1: attGAAAAGGGCAAGGAGGTGGTgttttagagctagaaatagcaag,
sgRNA_3: attGGCGGAGGAGCAGCCGCAAGgttttagagctagaaatagcaag,
sgRNA_2: attGGGCGGACTCACGTTAGTCAgttttagagctagaaatagcaag,
sgRNA_4: attGTTGGAGCGCGTGAGGATCAgttttagagctagaaatagcaag.
Regions that map to the locus are in capital letters, mapping immediately upstream of the PAM motif (NGG). The long 3'-tail corresponds to the scaffold for Cas9 binding. Two *FLClean* lines were created *FLClean_13*, with sgRNA 1 and 3, and *FLClean_24,* with sgRNA 2 and 4. The *FLClean_24* line was used as background for floral dipping with the *TEX1.0* and *TEX2.0* constructs. The *pVRN5:VIN3-eGFP* line was generated by replacing the *VIN3* promoter sequence from the *pVIN3:VIN3-eGFP* construct described previously (5) with the promoter sequence of *VRN5*. The construct was transformed into *vin3-4 FRI* by floral dipping, and individual lines were selected. Two individual lines (#28 and #40) were used for ChIP analysis.

#### Growth conditions

Seeds were surface sterilized and sown on 1x Murashige and Skoog (MS) media without glucose. As *FLC* shutdown in cold is sensitive to growth, seeds for expression analysis were sown at low density. Seeds were stratified for 2–3 days at 4ºC and grown for 10 days under long-day conditions (16h light, 8h dark at 20ºC). For vernalization treatment, seedlings were transferred to short-day conditions (8h light, 16h darkness at constant 5ºC) after 10 days pre-growth. Plants harvested 10 days after vernalization were transferred back to long-day conditions and harvested from plates. For longer post-vernalization treatment (more than 10 days), the plants were transferred to soil and grown under long-day conditions. Fluctuating cold conditions used in Fig. 3 were as described previously (1).

**Expression analysis**

Total RNA was extracted using the hot phenol method as described previously (6). Genomic DNA contamination was removed with TURBO DNase (Invitrogen), following the manufacturer's guidelines. cDNA was synthesized with SuperScript IV reverse transcriptase (Invitrogen). Gene-specific primers were used for reverse transcription (RT) of *COOLAIR*, *FLC*, and *VIN3*. For the RT reaction to analyse the expression of vernalization factors in SI Fig. 1, oligo(dT) primers were used. Quantitative PCR (qPCR) was performed using SYBR Green I Master (Roche) and analysed on a LightCyler 480 machine (Roche). Ct values were normalized to the geometric mean of *UBIQUITIN CARRIER PROTEIN 1 (UBC)* and *SERINE/THREONINE PROTEIN PHOSPHATASE 2 A (PP2A)*. All primers are listed in SI Appendix, Table S1.

**Chromatin immunoprecipitation**

Chromatin immunoprecipitation (ChIP) was performed as described in (7). ChIP was performed with antibodies: α-H3 (Abcam, ab1791), α-H3K36me3 (Abcam, ab9050), α-H3K27me3 (Abcam, ab192985), α-H3K27me2 (Upstate, 07-452), α-H2AK119ub (Cell Signaling Technology, #8240), α-H3K4me1 (Abcam, ab8895), and α-GFP (Abcam, ab290). ChIP enrichment was quantified by qPCR with primers listed in SI Appendix, Table S1. For histone, ChIP values were normalized to H3. For H3K36me3 and H3K27me3, values were further normalized to enrichment at the positive control loci *ACTIN* for H3K36me3 and *STM* for H3K27me3. Primers; *FLC* 0, *FLC* 250, and *FLC* 500 were used to calculate the average level in the nucleation region for the plots in Fig. 2 and SI Fig. 5.

**Chromatin Conformation Capture**

Chromatin conformation capture was performed as described previously (8, 9) with minor modifications. 1 g of 10 days old seedlings were crosslinked in 2 % formaldehyde for 20 mins. Crosslinking was stopped by the addition of 2M Glycin to a final concentration of 0.125 M and vacuum infiltrated for 7 mins. Nuclei were extracted with Honda buffer as for ChIP (7). After purification, chromatin was digested with 600 U of BamHI (NEB) and BglII (NEB) for 14-16 h, followed by 8h of ligation with T4 ligation (Promega) at 17°C. DNA was purified with Phenol:Chloroform:IAA (25:24:1) and precipitated with isopropanol. DNA was dissolved in water and further purified using the ChIP DNA Clean & Concentration (Zymo Research) kit, following the manufacturer's protocol. The 3C library was quantified as described previously (8).

### Mathematical modelling

In this study we use mathematical modelling to dissect the complex interplay of Polycomb mediated silencing and antisense mediated repression involved in *FLC* regulation in the cold. The models used in this study are constructed within a framework we have previously developed and experimentally validated. This framework describes the dynamics of transcriptional shutdown and histone modification changes at the whole-plant level during cold-induced epigenetic silencing at *FLC* (10, 11). Therefore, the assumptions used in these previous models are carried through into the models developed here. Many of these assumptions are directly based on experimentally established details, while the validity of others has been established through experimental testing of model predictions in these previous studies (10, 11). The models developed in this study use these assumptions as a starting point before we add additional features.

##### H3K36me3 and H3K27me3 dynamics in *COOLAIR* defective mutants

Our previously developed (and experimentally validated) models successfully captured the behaviour of H3K27me3 at *FLC* in cold and post-cold conditions observed in Col*FRI*, nucleation mutants, spreading mutants, as well as reactivation in a natural variant of *FLC*. These models also captured some, although not all, aspects of H3K36me3 dynamics. However, these models did not capture the role of antisense mediated regulation. Here, we start with the existing modelling framework, and attempt to build a model that, in addition to what is captured by previous models, can also properly incorporate the dynamics of H3K36me3 and H3K27me3 at *FLC* as observed in Col*FRI* and the *COOLAIR* defective mutants. To do this we focus on only those trends which are consistent across the three different *COOLAIR* defective mutants. These features are as follows (Fig. 2):

1. Similar dynamics of H3K27me3 nucleation and spreading in Col*FRI* and *COOLAIR* defective mutants, with a significant difference in starting (NV) levels.
    - H3K27me3 nucleation is not slower in *COOLAIR* defective mutants.
    - H3K27me3 levels at 6WT0 are similar in Col*FRI* and *COOLAIR* defective mutants.
    - H3K27me3 spreading is similar in Col*FRI* and *COOLAIR* defective mutants.
2. Significant differences in H3K36me3 dynamics of the *COOLAIR* defective mutants during the cold.
    - H3K36me3 increases across the locus (relative to NV) in the *COOLAIR* defective mutants in the first two weeks of cold, while Col*FRI* does not show this trend.

- The overall reduction in H3K36me3 levels over six weeks of cold is clearly weakened in the *COOLAIR* defective mutants, so that they exhibit significantly higher levels of this modification across the locus at 6WT0.
- Consistent with H3K36me3 changes and the association of this modification with transcription, *FLC* unspliced also reduces more slowly in the *COOLAIR* defective mutants.

3. Similar behaviour of H3K36me3 in *COOLAIR* defective mutants and in Col*FRI* during post-cold growth.
    - A clear reduction of H3K36me3 levels is observed between 6WT0 and 6WT10/T20, in both *COOLAIR* defective mutants and Col*FRI.*
    - Consistent with H3K36me3 reduction, and the association of this modification with transcription, *FLC* transcriptional output also reduces further in the post-cold.

**Building a model to capture the differences in H3K36me3 dynamics**

The behaviour of H3K27me3 in the *COOLAIR* defective mutants (described above) can be captured by previous models, since the behaviour is similar to Col*FRI*, except for the difference in NV levels. However, these models cannot capture the H3K36me3 behaviour, which shows significant differences in the *COOLAIR* defective mutants. This is because: (i) these models do not explicitly include a regulatory role for antisense transcription, and (ii) these models treat H3K36me3 and H3K27me3 as being exclusively present in different states of the *FLC* locus (H3K36me3 only in a high transcriptional state, and H3K27me3 only in a nucleated or spread state).

**Capturing *FLC* states at the whole plant level**

The existing modelling framework describes *FLC* locus states in a population of cells representing the whole plant. This population is made up of dividing and non-dividing cells (10,11). One of the core assumptions of these models is that the *FLC* locus can undergo nucleation of H3K27me3 in both dividing (meristematic tissue) and non-dividing cells. The validity of this assumption is supported by both direct and indirect experimental evidence: (i) for dividing cells, measurements of *FLC* epigenetic silencing by Polycomb in root meristematic cells via fluorescent imaging of FLC-Venus in plants defective for spreading of H3K27me3 (12); (ii) for non-dividing cells, the direct measurement of H3K27me3 nucleation by ChIP in mature leaves during vernalization in Col*FRI* (13); (iii) for non-dividing cells, the indirect evidence from all of our own ChIP time course datasets for H3K27me3: If only copies in dividing cells were capable of nucleation, the repeated division of these nucleated copies during rapid post-cold growth would cause a significant increase in population-level nucleation region H3K27me3. The fact that no such increase is observed by ChIP during post-cold growth indicates that *FLC* copies in both dividing and non-dividing cells can nucleate. This continues to be a core assumption in the models built here. As discussed in the main text, our data indicates that the antisense mediated repression and the PRC2/H3K27me3 mediated silencing function in parallel at *FLC*. This paradigm of two parallel pathways is therefore also central to the models developed here.

**Possible models with different behaviour in subpopulations of cells**

With the previously observed mutual exclusivity of H3K27me3 and H3K36me3 at *FLC* during vernalization (14), as well as other evidence for the mutual exclusivity of these modifications (15), it is tempting to consider models where apparent disruption of this mutual exclusivity in *COOLAIR* defective mutants arises from different subpopulations of *FLC* copies ending up in different states. While some tissue specific behaviour cannot be ruled out, here we examine two simple models with different behaviour between subpopulations and demonstrate that such subpopulation specific behaviour by itself is insufficient to capture the observed trends.

**Model 1:** A simple approach to explain the difference in H3K36me3 dynamics during the cold between Col*FRI* and the *COOLAIR* defective lines, using the existing models, would be to introduce a subpopulation of cells *only* in the *COOLAIR* defective mutants, in which H3K27me3 does not nucleate at *FLC* in the cold. These *FLC* copies would remain in an active transcriptional state, thus producing a higher level of H3K36me3 during the cold in the *COOLAIR* defective lines. However, the presence of such a non-nucleating subpopulation would be expected to produce a clear reduction of nucleation region H3K27me3 levels (compared to Col*FRI*) during the cold. Since this is not observed, particularly at 6WT0, we reject this model.

**Model 2:** A more sophisticated model to explain the difference in H3K36me3 dynamics in the cold is one where we again have a subpopulation of non-nucleating cells, but this subpopulation is common to both Col*FRI* and the *COOLAIR* defective mutants. This subpopulation has to include roughly the same proportion of dividing cells as the rest of the population (otherwise this model would produce a post-cold increase in nucleation region H3K27me3, which as discussed above, is inconsistent with all our data). The existence of such a subpopulation would allow H3K27me3 dynamics to be unchanged between these genotypes. The higher H3K36me3 in the *COOLAIR* defective mutants could then be explained by antisense mediated repression having a role specifically in the non-nucleating subpopulation. However, since the non-nucleating population includes dividing cells, their active *FLC* states (and hence high H3K36me3 levels) would be maintained and propagated during post cold growth. Therefore, without invoking additional, unknown mechanisms for H3K36me3 removal in this subpopulation of cells, even this model cannot explain the post cold reduction in H3K36me3 leading to essentially the same levels in Col*FRI* and the *COOLAIR* defective mutants by 6WT10.

Thus, we are led to construct a model where H3K27me3 and H3K36me3 can co-exist at the same *FLC* copy, and where antisense transcription modulates H3K36me3 levels. We note that this coexistence of the two marks could potentially even involve coexistence on the same H3 tail - H3K27me3 accumulation at *FLC* during cold is mediated by a VRN2-PRC2 complex, whose activity is insensitive to the presence of H3K36me3 on a substrate H3 tail (15). Our mathematical model is detailed below.

#### Model features

**Chromatin states at *FLC* copies:** The model allows each copy to be in one of three states: an active transcriptional state (no H3K27me3 nucleation), an H3K27me3 nucleated state, and an H3K27me3 spread state. Importantly, the model also allows H3K36me3 to be present at the locus in *all of these states*, at a level that is determined by the transcriptional activity possible in each state rather than direct mutual exclusivity with H3K27me3. This means that the *FLC* copies in the active transcriptional state (no H3K27me3 nucleation) make the highest contribution to population level H3K36me3 levels; copies in the H3K27me3 nucleated state have lower transcriptional activity and hence make an intermediate contribution to population level H3K36me3; copies in the H3K27me3 spread state have the lowest transcriptional activity and hence make the lowest contribution to population level H3K36me3.
We note that the above features replace the assumptions used in previous models (10,11) :

(1) Allowing H3K36me3 to be present in all states at a level determined by transcriptional activity replaces the previous assumption that this modification is only present in a high transcriptional state of *FLC.*
(2) Having only three states of the locus – active, H3K27me3 nucleated, and H3K27me3 spread – replaces the assumption that there is a distinct, “inactive” state of the locus with neither H3K36me3 nor H3K27me3 accumulation, which is set up by a “VIN3 independent” pathway.

The new assumptions allow the model to capture transcriptional downregulation and H3K36me3 levels in parallel to H3K27me3 mediated changes of transcriptional state, and thus emphasises the paradigm that emerges from our data – that of parallel pathways (antisense transcription mediated and PRC2 mediated) converging to regulate *FLC* expression.

**Dividing and Non-dividing loci:** The total number of dividing copies is fixed**.** We use a simplified division model (11), where each division produces one dividing copy and a fixed number of non-dividing copies.

**Nucleation and Spreading:**

- Transitions from an active to a nucleated state are allowed only in the cold, with the probability of nucleation dictated by the VIN3 protein concentration calculated from the LSCD model of VIN3 dynamics (10) – a predictive model of VIN3 expression that captures the effect of multiple thermosensory inputs operating at different timescales.
- The transition from a nucleated to a spread state occurs during replication/division, consistent with the dependence of spreading on an active cell cycle that we have previously shown (12).
- Replication/division causes a transition from a nucleated to a spread state: each division of a nucleated copy produces one dividing, spread copy and a fixed number of non-dividing, spread copies.
- The spread state is assumed to be stable – the model does not allow reactivation from the spread state.
- Except in simulations of the spreading mutant, we assume no loss of nucleation at nucleated, dividing copies.

**Division rate and pre-growth duration**

- The growth rate is assumed to undergo step change in cold (reduced by a factor of 40 in cold) (11).
- The pre-growth duration is fixed at 10 days (11).

**Sense transcriptional activity (initiation rate):** Antisense mediated regulation of sense transcription (initiation rate) is assumed to be possible in all three states – active, nucleated, and spread. The highest level of transcriptional activity in the nucleated state assumes 0.3 of the highest level in the active state. Based on our data, which shows that cold induced H3K27me3 nucleation and post-cold H3K27me3 spreading are not disrupted in the *COOLAIR* defective mutants, we assume that nucleation and spreading of H3K27me3 (i.e. the rates of transition to these states) are unaffected by the level of transcriptional activity.

**Sense transcriptional activity (elongation rate):** The increase in H3K36me3 between NV and 2WT0 observed in the *COOLAIR* defective mutants cannot be captured by only having co-transcriptional addition of this modification – there is a general trend towards reduction rather than an increase in sense transcriptional activity between these timepoints (as measured by *FLC* unspliced and spliced transcript levels). Therefore, to capture this increase in H3K36me3, we introduce a reduction in the elongation rate in the cold: it is assumed to drop to 0.6 of its NV value in the cold and also recovers post-cold. Such a reduction in elongation rate allows the Pol II density at the locus to increase between the NV and 2WT0 timepoints, even with a reduction in transcriptional activity (initiation rate) between these timepoints.

**Antisense mediated (nucleation independent) repression pathway:** A nucleation independent repression pathway performs analogue control of the sense transcription level in active and nucleated states**.** The functioning of this pathway is assumed to rely on antisense (AS) transcription (i.e., this pathway is not functional in the *COOLAIR* defective mutants). Repression by this pathway is assumed to increase slowly in the cold and decrease quickly upon return to warm, consistent with NTL8 dynamics (16). This repression is modelled as a slow reduction in the transcription initiation rate in the cold, and a rapid recovery in the initiation rate in the post-cold. This is captured by a multiplicative factor set to vary between 1 and 0.5 with an exponential decay over time in cold and a faster exponential recovery over time in the warm (see model implementation below). This is consistent with our previous model-predicted NTL8 accumulation dynamics determined by slower growth in the cold (16), and rapid NTL8 reduction during post-cold growth, as well as the measured slow build-up of *FLC* antisense transcripts in constant cold measured by qPCR (17) This is also consistent with the analysis of the VIN3 independent pathway in (10) – the dynamics of this pathway was predicted to be temperature dependent, causing slow *FLC* reduction in the cold, but allowing rapid increase in the post cold in the absence of VIN3 dependent H3K27me3 nucleation. The reduction of the transcriptional initiation rate (caused by the AS pathway) is assumed to have the same dynamics at active (non-nucleated) and nucleated copies. The transcriptional initiation recovery timescale in the post-cold is also assumed to be the same at non-nucleated and nucleated copies.
We note that the above set of assumptions describing the antisense mediated pathway replaces population level H3K36me3 the simpler description of a “VIN3 independent” pathway used in our previous models (10,11).

**H3K27me3 levels:** The contribution to H3K27me3 levels from an individual *FLC* locus depends on the state of the locus: low in the nucleation region (NR) and non-nucleation region (body) for active copies, high in the NR and low in the body for nucleated copies, high in the NR and body for spread copies.

**H3K36me3 levels in the NR depends on Pol II density:** For simplicity, the model describes the H3K36me3 levels in the *FLC* nucleation region, but we note that the levels of this modification across the gene body follow the trends in the nucleation region in all our data (Fig. 2B, (12,14)), with the only exception being the 3’ end of the locus, where H3K36me3 levels reflect the level of antisense transcription (increasing during the cold (Fig. 3B,C), (14) and high in ntl8-D3 (Fig. 1B)). Based on the above evidence, the H3K36me3 level is assumed to be proportional to the Pol II density. This is consistent with co-transcriptional addition of this modification (18), as well as longer Pol II dwell time at a given location allowing a greater window of opportunity for adding this modification (19). Both Pol II density and H3K36me3 levels are assumed to be at quasi-steady state, and the Pol II density is computed as the ratio of the initiation rate to the elongation rate. As described above, the initiation rate is determined by two factors: the H3K27me3 state and the antisense mediated repression pathway.

#### Model implementation

Following the same approach as for our previous models (10,11), an ODE (Ordinary Differential Equation) model is constructed using the above assumptions. The model equations are shown below. The model is simulated using the ode15s solver in Matlab version 2017a.

**Model Variables:**

$f_{a,d}$ (fraction of active, dividing copies)

$f_{n,d}$ (fraction of nucleated, dividing copies)

$f_{s,d}$ (fraction of spread, dividing copies)

$f_{a,nd}$ (ratio of active, non-dividing copies to total number of dividing copies)

$f_{n,nd}$ (ratio of nucleated, non-dividing copies to total number of dividing copies)

$f_{s,nd}$ (ratio of spread, non-dividing copies to total number of dividing copies)

The dynamics are such that $f_{a,d} + f_{n,d} + f_{s,d} = 1$ at all timepoints.

**Basic Model (representing Col*FRI*):**

$$\frac{df_{a,d}}{dt} = -k_s f_{a,d}$$

$$\frac{df_{n,d}}{dt} = k_s f_{a,d} - g(T) f_{n,d}$$

$$\frac{df_{s,d}}{dt} = g(T) f_{n,d}$$

$$\frac{df_{a,nd}}{dt} = d_n g(T) f_{a,d} - k_s f_{a,nd}$$

$$\frac{df_{n,nd}}{dt} = k_s f_{a,nd}$$

$$\frac{df_{s,nd}}{dt} = d_n g(T) f_{n,d} + d_n g(T) f_{s,d}$$

Here, $g(T)$ represents the division rate, which undergoes a step change reduction in the cold. $k_s$ represents the nucleation rate, computed using the VIN3 level as in (11), where the VIN3 level is itself computed using the LSCD model (10). $d_n$ represents the number of non-dividing copies produced at each division event (using the same simplified description as in (11).

**Model for *COOLAIR* defective mutants:** The basic model is used with no changes. The difference is in processing the simulation output (see below).

**Model for H3K27me3 Nucleation mutants:** The basic model is used with the nucleation rate $k_s$ set to zero throughout the simulation. The initial conditions in this case are still allowed to have a non-zero fraction of H3K27me3 spread *FLC* copies, consistent with the NV level of H3K27me3 observed in cold-nucleation mutants including *vrn2-1* and *vin3-4* (12).

**Model for H3K27me3 Spreading mutant:** Here we modify the basic model to capture reactivation/loss of H3K27me3 nucleation. At each division event, a nucleated dividing copy is allowed to undergo three different scenarios (11):

- Reactivation, producing one active dividing copy and $d_n$ active non-dividing copies.
- Spreading, producing one spread dividing copy and $d_n$ spread non-dividing copies.
- Neither spreading nor reactivation, producing one nucleated dividing copy, $\beta_{nuc} d_n$ nucleated non-dividing copies, $\beta_{sprd} d_n$ spread non-dividing copies, and $\beta_{react} d_n$ active non-dividing copies.

$$\frac{df_{a,d}}{dt} = -k_s f_{a,d} + \gamma g(T) f_{n,d}$$

$$\frac{df_{n,d}}{dt} = k_s f_{a,d} - \delta g(T) f_{n,d} - \gamma g(T) f_{n,d}$$

$$\frac{df_{s,d}}{dt} = \delta g(T) f_{n,d}$$

$$\frac{df_{a,nd}}{dt} = d_n g(t) f_{a,d} - k_s f_{a,nd} + d_n \gamma g(T) f_{n,d} + \beta_{react} d_n (1 - \delta - \gamma) g(T) f_{n,d}$$

$$\frac{df_{n,nd}}{dt} = k_s f_{a,nd} + \beta_{nuc} d_n (1 - \delta - \gamma) g(T) f_{n,d}$$

$$\frac{df_{s,nd}}{dt} = d_n \delta g(T) f_{n,d} + d_n g(T) f_{s,d} + \beta_{sprd} d_n (1 - \delta - \gamma) g(T) f_{n,d}$$

Here $\gamma$ represents the fraction of nucleated dividing copies undergoing reactivation and $\delta$ represents the fraction of nucleated dividing copies undergoing spreading, at each replication/division event. The parameters $\beta_{nuc}$, $\beta_{sprd}$, and $\beta_{react}$ were computed numerically (see SI table 1 of parameter values), using a Monte-Carlo approach to carry out five successive replication/division events starting from one nucleated copy (five divisions is consistent with our assumption of $d_n = 32$ (11).

**Processing simulation output:**

The total fractions of copies in each state (at any timepoint) can be computed as follows from the simulation output:

Total fraction of active copies: $F_a = \dfrac{f_{a,d} + f_{a,nd}}{1 + f_{a,nd} + f_{n,nd} + f_{s,nd}}$

Total fraction of nucleated copies: $F_n = \dfrac{f_{n,d} + f_{n,nd}}{1 + f_{a,nd} + f_{n,nd} + f_{s,nd}}$

Total fraction of spread copies: $F_s = \dfrac{f_{s,d} + f_{s,nd}}{1 + f_{a,nd} + f_{n,nd} + f_{s,nd}}$

The total fractions of copies in each state are then used to compute the histone modification levels as follows:

NR H3K36me3 level: $K36_{NR} = p_{K36}\left(\dfrac{q(t)\left(r_a F_a + r_n F_n\right) + r_s F_s}{v(T)}\right)$

NR H3K27me3 level: $K27_{NR} = p_{K27}\left((0)F_a + (1)F_n + (1)F_s\right)$

Body H3K27me3 level: $K27_{Body} = p_{K27}\left((0)F_a + (0)F_n + (1)F_s\right)$

The multiplicative factors $r_a$ , $r_n$ and $r_s$ represent the maximum transcription initiation rate for active, nucleated, and spread copies respectively. $v(T)$ represents the Pol II elongation rate, which is assumed to undergo a step change (reduction) during the cold. The parameters $p_{K36}$ and $p_{K27}$ are used for scaling the model output for comparison to ChIP data. Note that having the parameter $r_n > 0$ means that copies in an H3K27me3 nucleated state can also contribute to the H3K36me3 levels calculated by the model. This reflects the model assumption that these two modifications can coexist in the nucleation region at a single *FLC* copy during cold induced silencing, with the H3K36me3 levels being limited only by the level of sense transcription. This assumption is based on the ability of the VRN2-PRC2 complex (which mediates cold induced H3K27me3 accumulation at *FLC*) to methylate H3 histones even when they carry K36 methylation (15).

The time-dependent multiplicative factor $q(t)$ capture the repression by the antisense mediated pathway. This factor has a basal value of 1 and decays exponentially with time to 0.5 in the cold and recover exponentially with time in the post-cold.

For simulating the *COOLAIR* defective mutants, the antisense mediated pathway is assumed to be non-functional. This is captured by setting $q(t) = 1$ throughout the simulation.

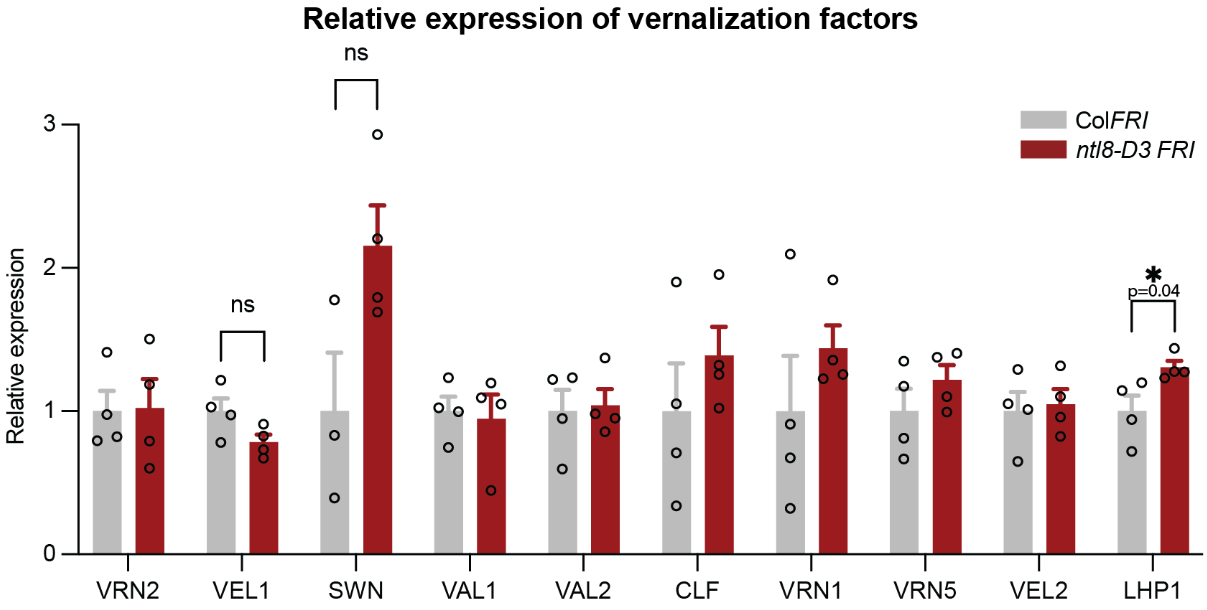


**Fig. S1. Relative expression of other factors involved in vernalization.** Expression of vernalization factors in *ntl8-D3 FRI* compared to Col*FRI* in non-vernalized conditions. Data are presented as the mean ± s.e.m ($n \geq 3$). Asterisks indicate significant different ($p \leq 0.05$, two-tailed t test). n.s, not significant. Each open circle represents a biological replicate.

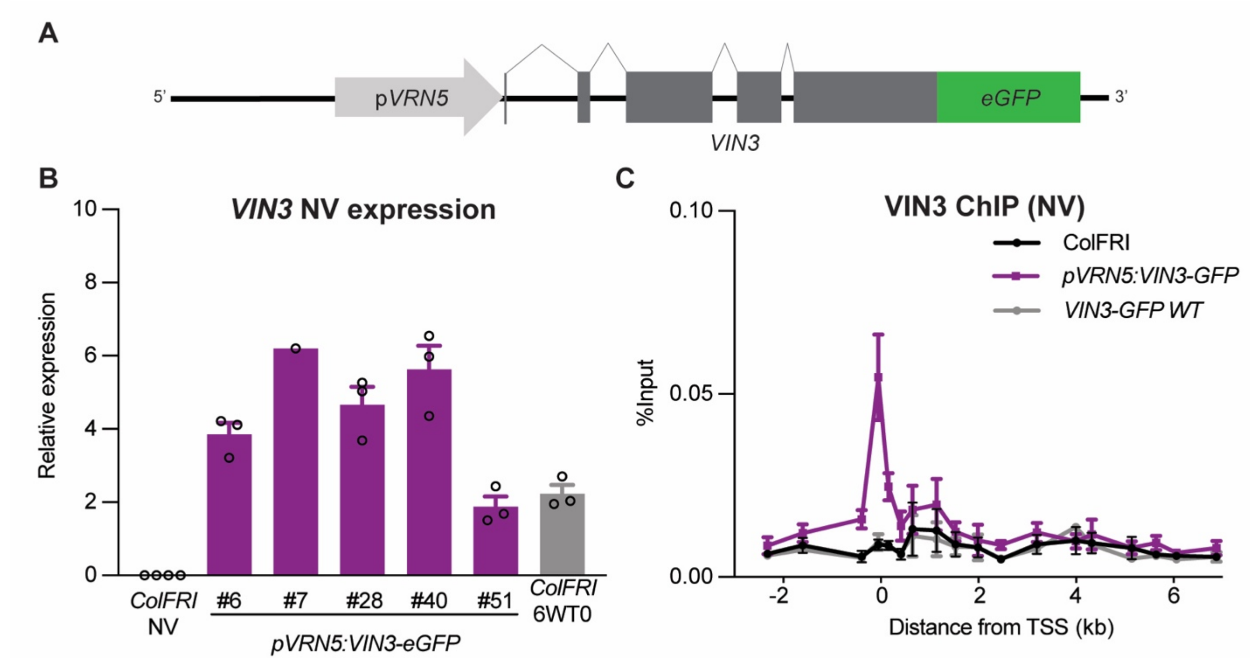


**Fig. S2. Constitutive expressed VIN3 binds the nucleation region at *FLC*. (A)** Schematic of the construct used to generate transgenic lines that express VIN3 in the absence of cold. The *VIN3* promoter was exchanged with the promoter sequence of *VRN5* (*pVRN5*). (**B**) Expression of *VIN3* in non-vernalized conditions (NV), *VIN3* expression in Col*FRI* after six weeks of vernalization (6WT0) is shown for comparison. Data are presented as the mean ± s.e.m relative to the geometric mean of *PP2A* and *ACTIN*. Each open circle represents a biological replicate. The numbers under the bars refer to individual transgenic lines. (**C**) VIN3-eGFP ChIP-qPCR enrichment at *FLC* at NV. Data are shown as percentage input. Non-transgenic Col*FRI* plants were used as a negative control sample. Error bars represent mean ± s.e.m. (n = 3 biological replicates).

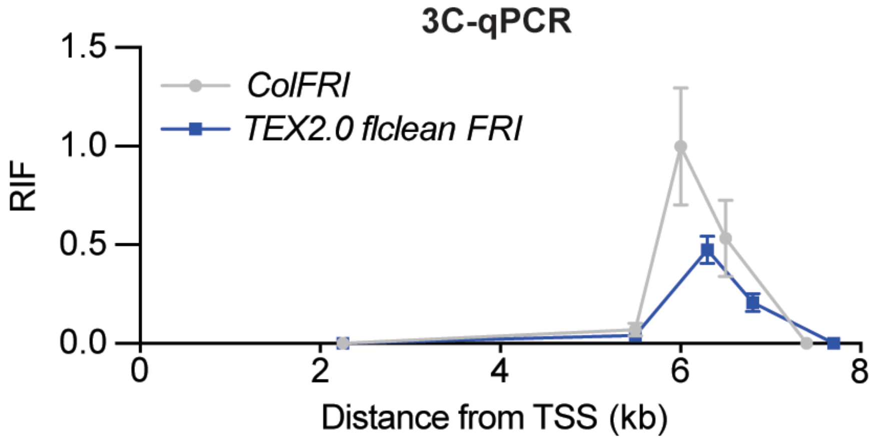


**Fig. S3. Gene-loop is disrupted in *TEX2.0*.** Quantitative 3C over the *FLC* locus in 10-day-old Col*FRI* and *TEX2.0 flclean FRI* with BamHI and BglII (similar to Fig. 1G). The region around the *FLC* transcription start site was used as the anchor region in the 3C analysis The data shows the relative interaction frequencies (RIF) and are the average of at least seven biological replicates. Data are presented as the mean ± s.e.m. (n ≥ 7).

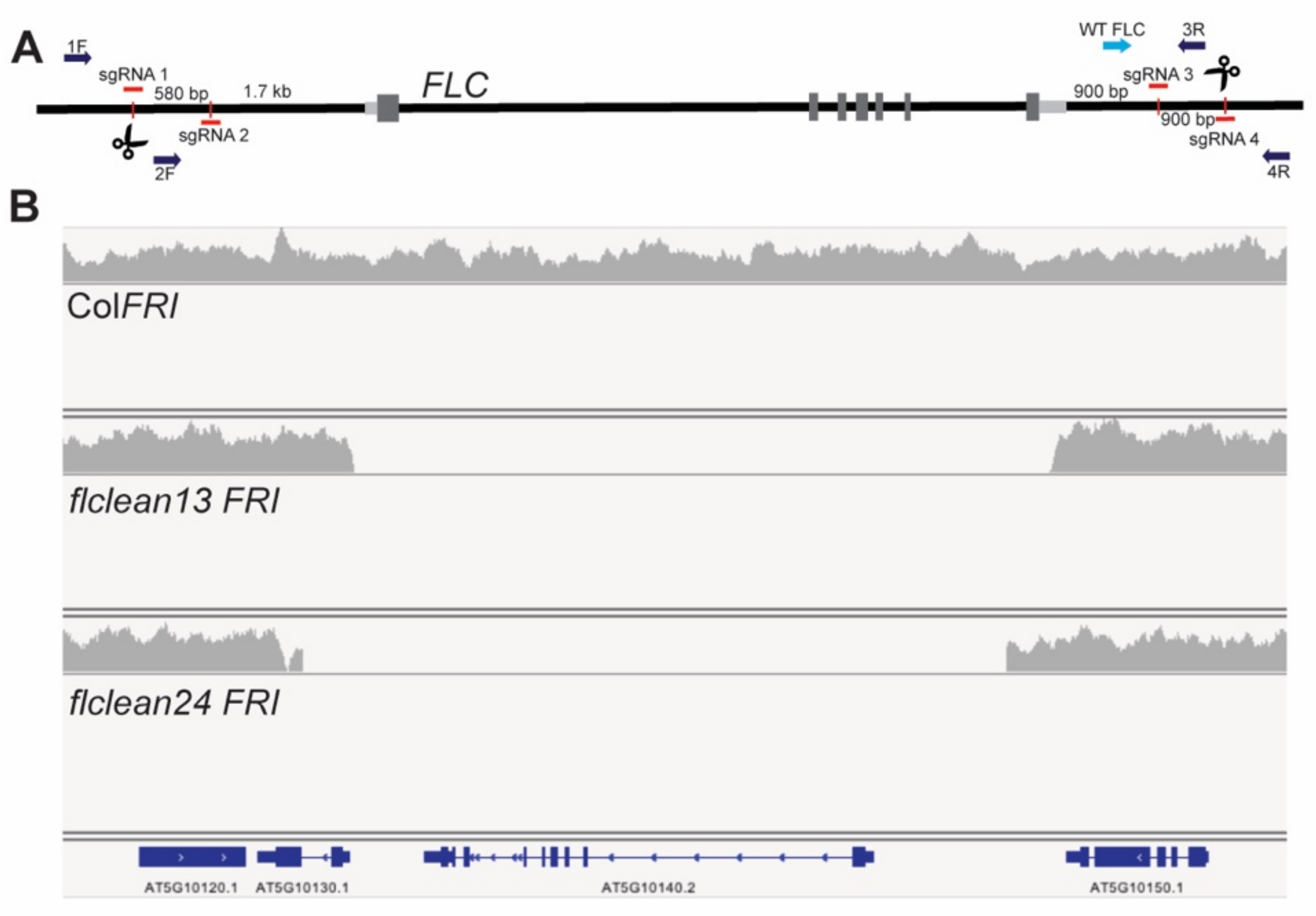


**Fig. S4. CRISPR/Cas9 mediated generation of *FLC* deletion lines (*FLClean*). (A)** Schematic representation of the *FLC* locus with locations of sgRNA target sites (Red) and primer binding sites for genotyping (Blue). The size numbers refer to the region at 5' and 3' end of *FLC* removed in the *FLClean* lines. **(B)** Integrative Genomics Viewer (IGV) screenshot of *FLC* genomic region showing read coverage of whole genome DNA sequencing in Col*FRI* and two CRISPR *FLClean* lines. The *FLClean* lines were created through removal of the whole *FLC* locus with either sgRNA1 and 3 (*FLClean13*) or sgRNA 2 and 4 (*FLClean24*).

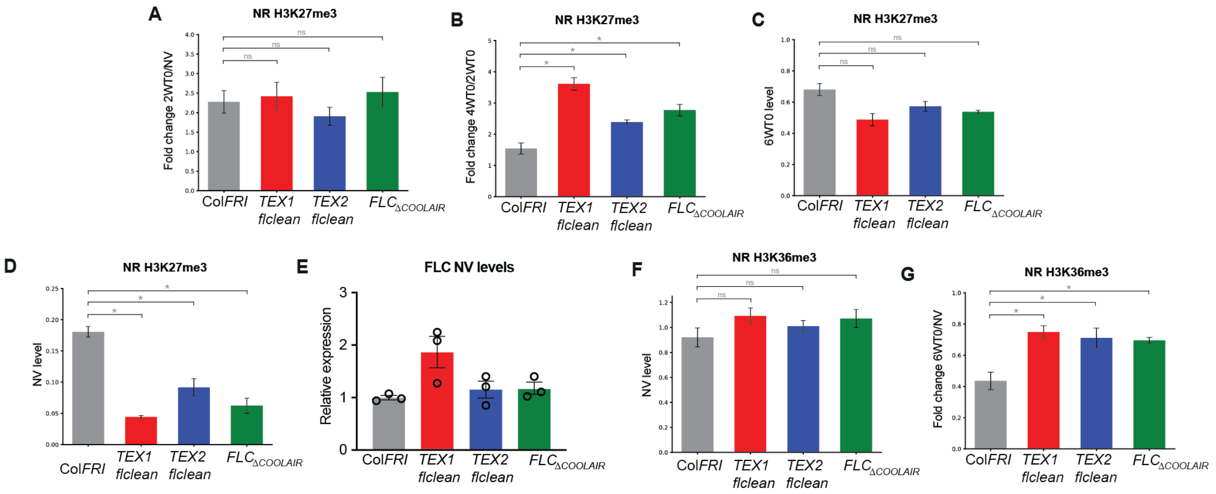


**Fig. S5. Quantitative analysis of antisense role in histone modification dynamics.** All comparisons shown consist of comparing the mean levels over three nucleation region primers between Col*FRI* to each of the defective *COOLAIR* lines (See supplementary information for details of primers). In cases where the qualitative trends were clear and consistent across the three defective *COOLAIR* lines, a one-tailed Student's t-test was used for each comparison. Error bars represent s.e.m. (n = 3 biological replicates). In (A), where there was no clear trend, a two-tailed Student's t-test was used. In all cases, the Bonferroni correction was used to adjust the significance level from $\alpha = 0.05$ to $\alpha = 0.0167$ (for three comparisons). (*) indicates $P < 0.0167$ ; ns indicates no significance ($P \geq 0.0167$). (**A**) Fold change (increase) of H3K27me3 in the nucleation region during first 2 weeks of cold treatment (2WT0/NV) is not significantly different between Col*FRI* and the defective *COOLAIR* lines. (**B**) Fold change (increase) of H3K27me3 in the nucleation region during the 2WT0 to 4WT0 period is significantly higher in the *COOLAIR* lines. (**C**) H3K27me3 levels at 6WT0 are not significantly lower in the defective *COOLAIR* lines. (**D**) NV level of H3K27me3 in the nucleation region is significantly higher in Col*FRI*. (**E)** *FLC* expression in 10 days old seedlings before cold exposure in Col*FRI* and the three defective *COOLAIR* lines; *TEX1, TEX2, and* $FLC_{\Delta COOLAIR}$*.* Unspliced RNA was measured and is shown relative to *UBC* and Col*FRI*. Error bars represent s.e.m. (n = 3 biological replicates). **(F-G)** Similar analysis as in (A-D). NV level of H3K36me3 is not significantly higher in the defective *COOLAIR* lines **(F)**. The fold change in H3K36me3 over 6W of cold treatment indicates significantly smaller changes in H3K36me3 in the defective *COOLAIR* lines **(G)**.

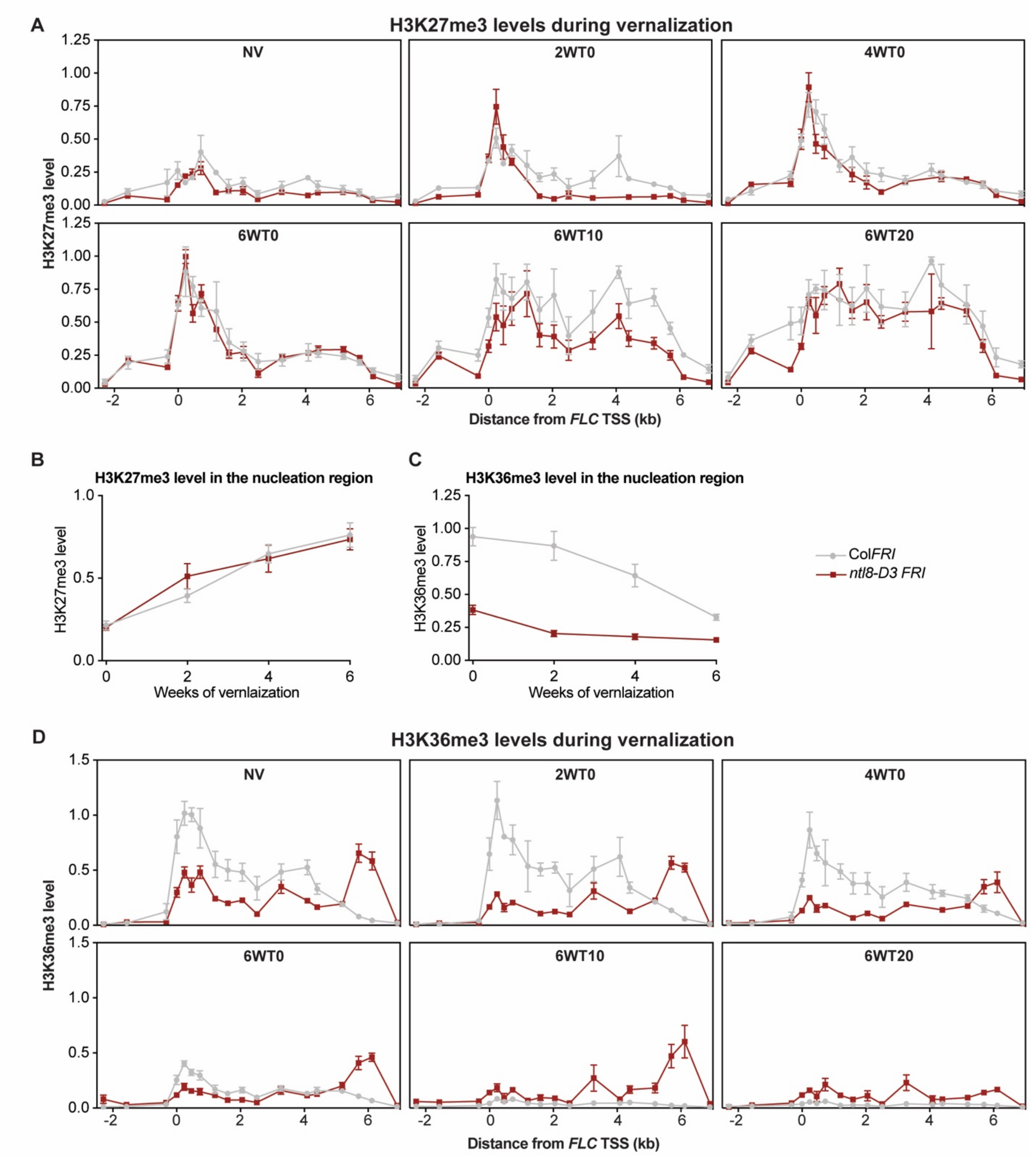


**Fig. S6. Polycomb repression of *FLC* is not inhibited by hyperactive antisense pathway in *ntl8-D3*. (A)** H3K27me3 ChIP in Col*FRI* and *ntl8-D3 FRI* across the *FLC* locus before, during, and after vernalization. H3K27me3 levels are expressed as relative to H3 and to the levels at the positive control gene *STM*. Data are presented as the mean ± s.e.m. (n ≥ 3). **(B)** H3K27me3 levels in the nucleation region in Col*FRI* and *ntl8-D3 FRI* during vernalization. The levels were calculated by averaging over three primers in the *FLC* nucleation region. **(C)** H3K36me3 levels in the nucleation region in Col*FRI* and *ntl8-D3 FRI* during vernalization. The levels were calculated as in B. **(D)** H3K36me3 ChIP in Col*FRI* and *ntl8-D3 FRI* across the *FLC* locus before, during, and after vernalization. H3K36me3 levels are expressed as relative to H3 and to the levels at the positive control gene *Actin*. Data are presented as the mean ± s.e.m. (n ≥ 3).

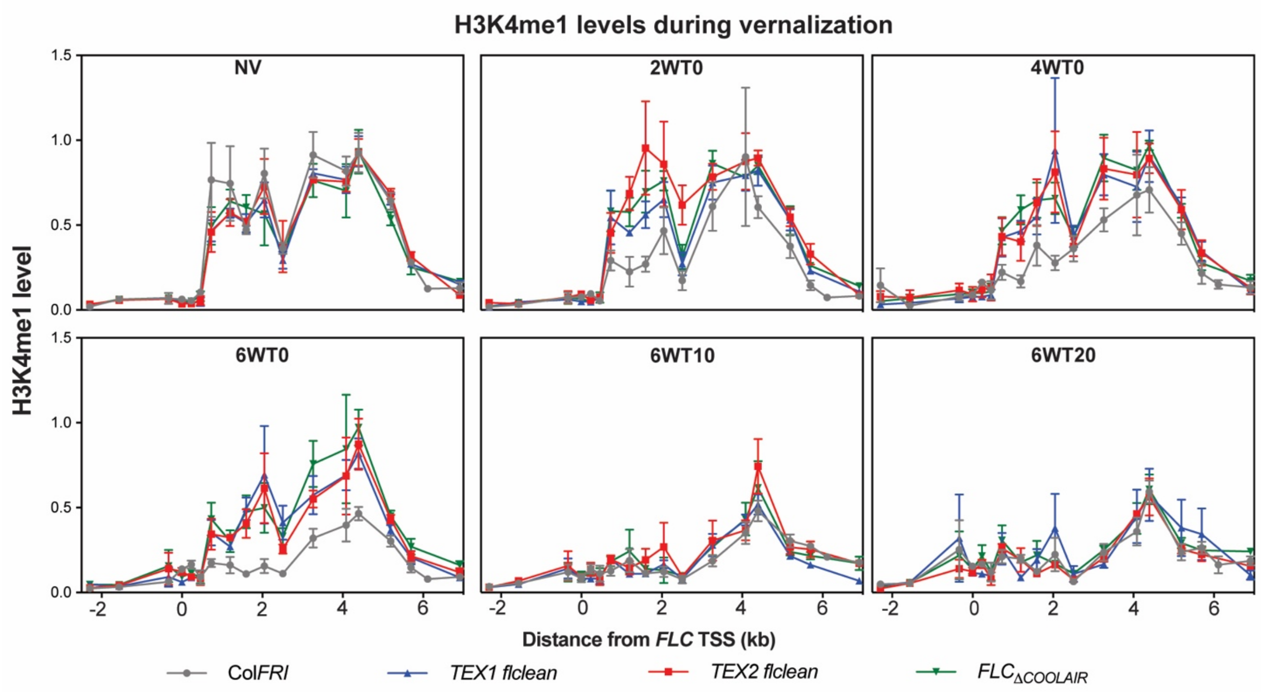


**Fig. S7. H3K4me1 removal during the cold is attenuated in COOLAIR defective lines.** H3K4me1 ChIP in Col*FRI* and the three COOLAIR mutant lines; *TEX1, TEX2, and* $FLC_{\Delta COOLAIR}$ across the *FLC* locus before, during, and after vernalization. Data are presented as the mean ± s.e.m. (n ≥ 3).

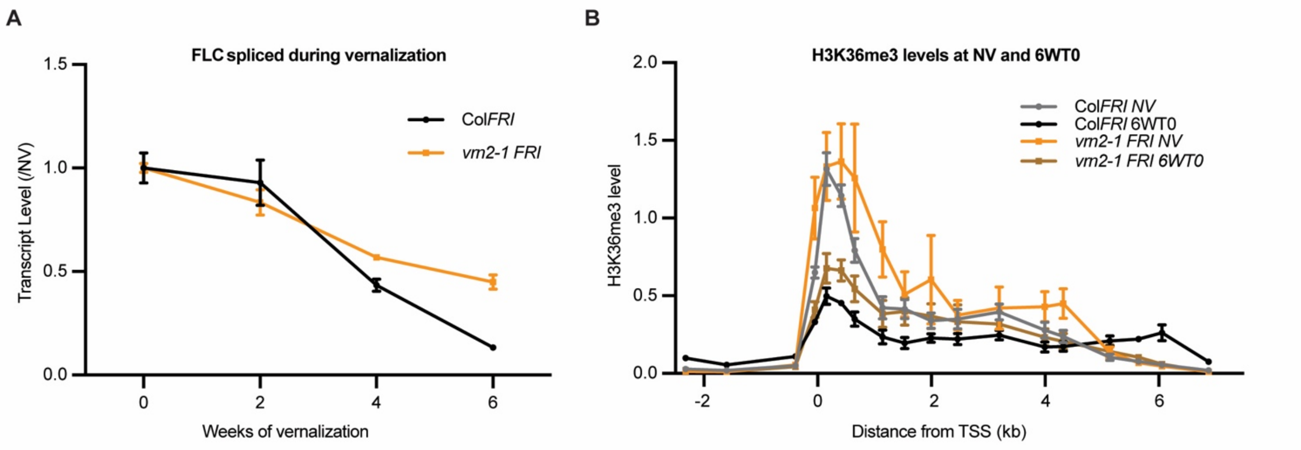


**Fig. S8. *FLC* repression is only partially disrupted in *vrn2-1 FRI*. (A)** *FLC* spliced levels and (B) H3K36me3 levels during vernalization in Col*FRI* and *vrn2-1 FRI*. **(A)** *FLC* spliced data shown relative to RNA levels in non-vernalized conditions (0 weeks of vernalization). **(B)** ChIP-qPCR data for H3K36me3 is normalized to H3 and shown relative to H3 normalized H3K36me3 levels at *ACTIN*. Data are presented as the mean ± s.e.m. ($n \geq 3$). Reproduced from (12).

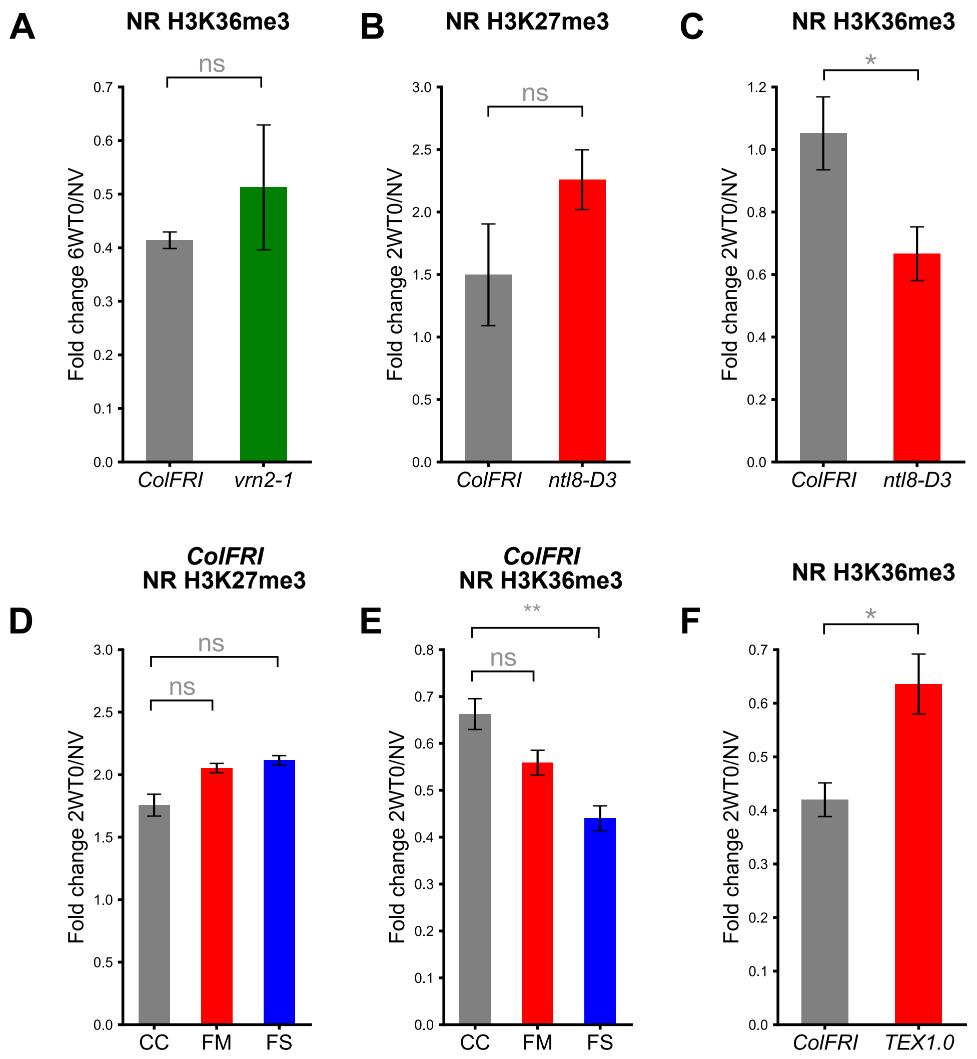


**Fig. S9. Fold change comparisons for changes in nucleation region H3K36me3 and H3K27me3.** This analysis is based on ChIP-qPCR time course data presented in Figs. 1,2. Error bars represent mean ± s.e.m. (n ≥ 3). All comparisons shown consist of comparing fold changes in the mean levels over three nucleation region primers in the indicated genotypes between different periods of cold and non-vernalized conditions (see supplementary information for details of primers). In all cases, the data showed clear trends, so a one-tailed Student's t-test was used for each comparison. In **(A-C)** and **(F)**, a significance level of alpha = 0.05 was used. (*) indicates P<0.05; ns indicates no significance (P≥0.05). In **(D)**, the Bonferroni correction was used to adjust the significance level from alpha = 0.05 to alpha = 0.025 (for two comparisons). ns indicates no significance (P≥0.025). In **(E)**, the Bonferroni correction was used to adjust the significance level from alpha = 0.01 to alpha = 0.005 (for two comparisons). (**) indicates P<0.005; ns indicates no significance (P≥0.005). **(A)** Comparison of NR H3K36me3 fold changes after 6 weeks cold (6WT0) in *ColFRI* and *vrn2-1.* **(B)** Comparison of NR H3K27me3 fold changes after 2 weeks cold

(2WT0) in *ColFRI* and *ntl8-D3*. **(C)** Comparison of NR H3K36me3 fold changes after 2 weeks cold (2WT0) in *ColFRI* and *ntl8-D3*. **(D)** Comparison of NR H3K27me3 fold changes after 2 weeks cold (2WT0) in *ColFRI* under different cold conditions – constant cold (CC), fluctuating mild (FM), and fluctuating strong (FS). **(E)** Comparison of NR H3K36me3 fold changes after 2 weeks cold (2WT0) in *ColFRI* under different cold conditions – constant cold (CC), fluctuating mild (FM), and fluctuating strong (FS). **(F)** Comparison of NR H3K36me3 fold changes after 2 weeks (2WT0) under FS conditions in *ColFRI* and *TEX1.0*.

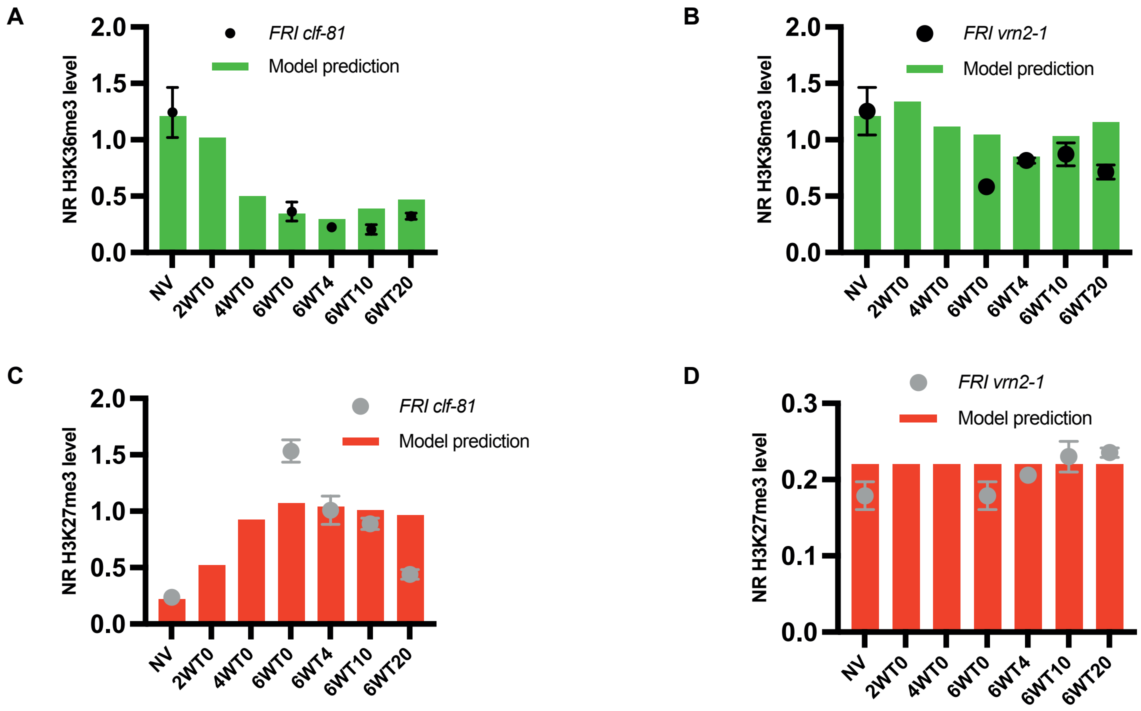


**Figure S10. Model predictions of impact of vernalization mutants on histone dynamics.** **(A-D)** Time course predictions from the mathematical model for other vernalization mutants: an H3K27me3 nucleation mutant, and a spreading mutant. The predictions are compared to previously published ChIP-qPCR time course data presented in (12).

Table S1. Model parameter values

| Parameter | Description | Value | Reference |
|---|---|---|---|
| $k_s$ | Rate constant of H3K27me3 nucleation ($day^{-1}$) | Computed as in (11), with $p_{s2} = 0.007\ \text{day}^{-1}{}^{\circ}\text{C}^{-2}$ | (10,11) |
| $g(T)$ | Temperature dependent growth rate ($day^{-1}$) | 0.4 for $T = 22$℃<br>0.01 for $T = 5$℃ | (11) |
| $d_n$ | Number of non-dividing copies produced per division of a dividing copy | 32 | (11) |
| $\delta$ | Fraction of H3K27me3 nucleated dividing copies undergoing spreading | 0.0025 | This allows for a very small fraction undergoing H3K27me3 spreading, even in a spreading mutant. Consistent with very low levels of H3K27me3 spreading observed in (12). |
| $\gamma$ | Fraction of H3K27me3 nucleated dividing copies undergoing reactivation during a replication/division event | 0.03 | Chosen to produce ~30% reactivation after 12 replication/division events starting from a single nucleated copy. This is based on the analysis of *FLC* reactivation in a spreading mutant in (12) |

| | | | |
|---|---|---|---|
| $\beta_{sprd}$ | Fraction of H3K27me3 spread non-dividing copies produced during replication of a nucleated copy in the spreading mutant model | 0.0121 | Numerically estimated from the same Monte Carlo simulation described above. |
| $\beta_{react}$ | Fraction of non-dividing copies that have lost H3K27me3 nucleation, produced during replication of a nucleated copy in the spreading mutant model | 0.1402 | Numerically estimated from the same Monte Carlo simulation described above. |
| $q(t)$ | Time-dependent multiplicative factor that captures repression by the antisense mediated pathway. | $q(t) = 1$ before cold<br>$q(t) = 0.5(1 + e^{-\alpha_1(t-10)})$ during cold (beginning at 10 days)<br>$q(t) = 1 - (1 - 0.5(1 + e^{-\alpha_1(42)}))e^{-\alpha_2(t-52)}$ during post-cold (beginning at 52 days) | This study |
| $\alpha_1$ | Rate constant of increasing antisense mediated repression during cold (day$^{-1}$) | 0.08 | This study |
| $\alpha_2$ | Rate constant of decreasing antisense mediated repression during post-cold (day$^{-1}$) | 0.12 | This study |
| $v(T)$ | Temperature dependent RNA Pol II elongation rate | 1 for $T = 22$℃<br>0.6 for $T = 5$℃ | This study |
| $r_a$ | Maximum transcription initiation rate for active copies (assumed normalised to rate at active copies) | 1 | This study |
| $r_n$ | Maximum transcription initiation rate for | 0.3 | This study |

| | | | |
|---|---|---|---|
| | nucleated copies (assumed normalised to rate at active copies) | | |
| $r_s$ | Maximum transcription initiation rate for spread copies (assumed normalised to rate at active copies) | 0.025 | This study |
| $p_{K36}$ | Scaling parameter for comparing model output to ChIP-qPCR data | 1.5 for *FRI clf-2 and FRI vrn2-1* (Data from (12))<br>1.3 for for *ColFRI*, and *COOLAIR* defective mutants (Data from this study) | This study |
| $p_{K27}$ | Scaling parameter for comparing model output to ChIP-qPCR data | 1.1 for *FRI clf-2 and FRI vrn2-1* (Data from (Yang et al., 2017))<br>0.7 for *ColFRI*, and *COOLAIR* defective mutants (Data from this study) | This study |

**Dataset S1**. List of primers used in this study
